# USP15: the fourth Proteasome-associated DUB

**DOI:** 10.1101/2025.08.13.670075

**Authors:** Shahar Levi, Fabian Glaser, Ajay R. Wagh, Eden Filimonov, Noa Reis, Sachin Mali, Sumeet Kumar Singh, Noam Cohen, Insuk Byun, Min Jae Lee, Oded Kleifeld, Ashraf Brik, Michael H Glickman, Indrajit Sahu

## Abstract

The human genome encodes approximately 100 deubiquitinating enzymes (DUBs), but only three are considered proteasome-associated DUBs (pDUBs): PSMD14/Rpn11, USP14, and UCHL5. Among these, only PSMD14 is an integral 19S subunit, whereas USP14 and UCHL5 bind transiently to specific proteasomal subunits. Given the dynamic nature of proteasome composition, we searched for additional pDUBs. USP15 was found to be associated with 26S proteasomes purified from cultured cells. In proteasome preparations from erythrocytes, USP15 was identified as the most abundant transient pDUB. It was even feasible to separate proteasomes containing USP15 from those containing USP14. Although USP15 utilizes an internal ubiquitin-like (UBL) domain for positioning itself at the proteasome, it did not compete with the UBL-containing USP14. USP15 facilitated substrate selection at the proteasome by efficiently disassembling short polyubiquitin (polyUb) chains, while sparing K48-linked tetra-ubiquitin conjugates from deubiquitination. This feature may aid the proteasome in differentiating between substrates to be rescued from those committed to proteolysis. Identification of a fourth pDUB encourages the continued search for additional proteasome-interacting proteins that modulate its substrate specificity in a context-specific manner.

## Introduction

The ubiquitin (Ub)-proteasome system (UPS) is an ATP-dependent pathway within eukaryotic cells that is responsible for breaking down misfolded, abnormal, and short-lived proteins (Finley, 2009; Schwartz & Ciechanover, 2009). This degradation process involves an enzymatic cascade that attaches Ub molecules to lysine residues on specific substrates, marking them for degradation by the proteasome. The prevalent signal for proteasomal degradation is K48-linked polyUb chains (Hershko & Ciechanover, 1998; van Tol et al., 2023). The proteasome holoenzyme is assembled from a cylindrical 20S core particle (CP) capped with one or two 19S regulatory particles (RPs), each consisting of the lid and base subcomplexes. The RP is made up of approximately 20 different integral subunits, as well as a handful of transiently associated proteasome-interacting proteins (PIPs) (Budenholzer et al., 2017; Sahu & Glickman, 2021; Verma et al., 2000). Some of the subunits have been ascribed functional roles in proteasome action. For instance, three subunits (PSMD2/Rpn1, PSMD4/Rpn10, and PSMD16/Rpn13) are Ub receptors that recognize and capture ubiquitinated substrates (Martinez-Fonts et al., 2020). Like other post-translational modifications, ubiquitination is reversible: proteases termed deubiquitinating enzymes (DUBs) can cleave the isopeptide bond between Ub and the substrate, edit Ub chains, and process Ub precursors (Komander et al., 2009).

The human proteome contains approximately 100 putative DUBs that can be separated into two main classes, cysteine proteases and metalloproteases (Harrigan et al., 2018; Komander et al., 2009). Of the cysteine-based DUBs, the Ubiquitin-specific proteases (USPs) form the largest known subfamily (~60) (Nijman et al., 2005). USPs are a subset of the papain superfamily and are characterized by their structural similarity and the extremely conserved catalytic triad: cysteine, aspartate/asparagine, and histidine residues (Barrett & Rawlings, 1996; Keijzer et al., 2024; Papa & Hochstrasser, 1993). USP family members differ from each other by additional sub-domains that add to their extensive structural and functional complexity by targeting them to intracellular compartments, substrates, or specific Ub linkages. For example, several DUBS – e.g. USP14, USP7, and USP15 – contain ubiquitin-like domains (UBL) which could target these proteins to Ub receptors on the proteasome (Hu et al., 2005; Nininahazwe et al., 2021), USP15, as well as USP11 and USP4, also harbors an autoregulatory domain present in USP (DUSP) domain.

Of the myriad DUBs, three are known to be associated with the 26S proteasome (pDUBs). One is the MPN^+^ metalloprotease (Maytal-Kivity et al., 2002), PSMD14/Rpn11, an integral 19S subunit, located in proximity to the polyUb-binding subunit PSMD4 (Sahu & Glickman, 2021; Yao & Cohen, 2002). Accessibility to the Ub receptor is crucial for its role in shaving the proximal Ub from substrates committed to degradation, and to relieve proteasome stalling by the globular Ub moiety (Greene et al., 2019; Htet et al., 2025; Verma et al., 2002; Yao & Cohen, 2002). Two other DUBs associate transiently with the proteasome: UCHL5/UCH37, which interacts with the PSMD16/Rpn13-PSMD1/Rpn2 unit (Hamazaki et al., 2006), and USP14, which associates with PSMD2/Rpn1 via its N-terminal UBL (Hu et al., 2005; Kim & Goldberg, 2018; Zhang et al., 2022). Specifically, UCHL5 has been demonstrated as a debranching DUB on the proteasome, removing ancillary branches from K48-linked Ub chains (Deol et al., 2020), whereas USP14 disassembles supernumerary K48-linked Ub chains from the substrate (Lee et al., 2016). In both cases, the enzymatic outcome of these pDUBs facilitates substrate processing post-engagement (Sahu & Glickman, 2021). Moreover, USP14 is prominently activated upon incorporation into the proteasome (Borodovsky et al., 2001; Hu et al., 2005; Lee et al., 2010, 2016; Zhang et al., 2022). The precise outcome can vary, since USP14 or its yeast ortholog Ubp6, have been shown to both stabilize cellular proteins against proteasomal degradation (Lee et al., 2010), or enhance proteasome proteolytic activity (Kim & Goldberg, 2018).

DUBs are one class of proteasome-interacting proteins (PIPs). The proteasome interactome is flexible, depending on the source, tissue, cell type, growth condition, or purification technique. Interestingly, erythrocytes and reticulocytes are abundant in UPS components, including proteasomes. Indeed, blood has been a common source of purified proteasome for enzymatic characterization (Matthews et al., 1989). Lately, proteasomes from numerous sources are available due to the ease of genetic manipulation in cell culture for affinity purification of tagged proteasomes (Byun et al., 2023). Knowing that proteasome composition is dynamic, and some pDUBs are transient, we searched for additional DUBs that may associate with proteasome complexes. Analysis of proteasome composition purified from erythrocytes in this study identified a subpopulation of 26S proteasome associated with USP15, but lacking USP14, essentially identifying a new population of proteasomes. This association was verified in different cell types by several methods, and we demonstrate that USP15 contributes to the sorting of ubiquitinated substrates committed to proteasomal degradation.

## Results

### USP15 found to be associated with purified human 26S proteasomes

To uncover potential DUB interactions, we performed an unbiased search for the presence of DUBs in different stages of a well-tested biochemical protocol for the purification of 26S proteasomes from erythrocytes (**Fig. 1A; Fig. Sup Fig. 1**). DUBs in each proteasome-containing sample were identified by MS/MS (**Spreadsheet 1**). Plotting the identified DUBs along the purification method, different patterns of behavior were discerned (**Fig. 1B**). Some DUBs, such as USP7, USP5, USP47, and USP4 diminished relative to proteasome subunits as the latter increased in purity (**Fig. 1B**; lower panel), suggesting that they are either unrelated to the proteasome or extremely labile. USP9x persistently showed up in proteasome fractions, yet the majority did not copurify with active 26S proteasomes throughout the purification steps. Considering its high MW (~300 KDa) it may be an unrelated copurifying protein that awaits future conformation. Interestingly, one DUB, USP15, maintained a consistent ratio with the integral proteasome subunits. In fact, it commigrated with the 26S proteasome in a similar fashion to UCHL5 and USP14, the two well-established pDUBs (**Fig. 1B**).

**Figure 1:**
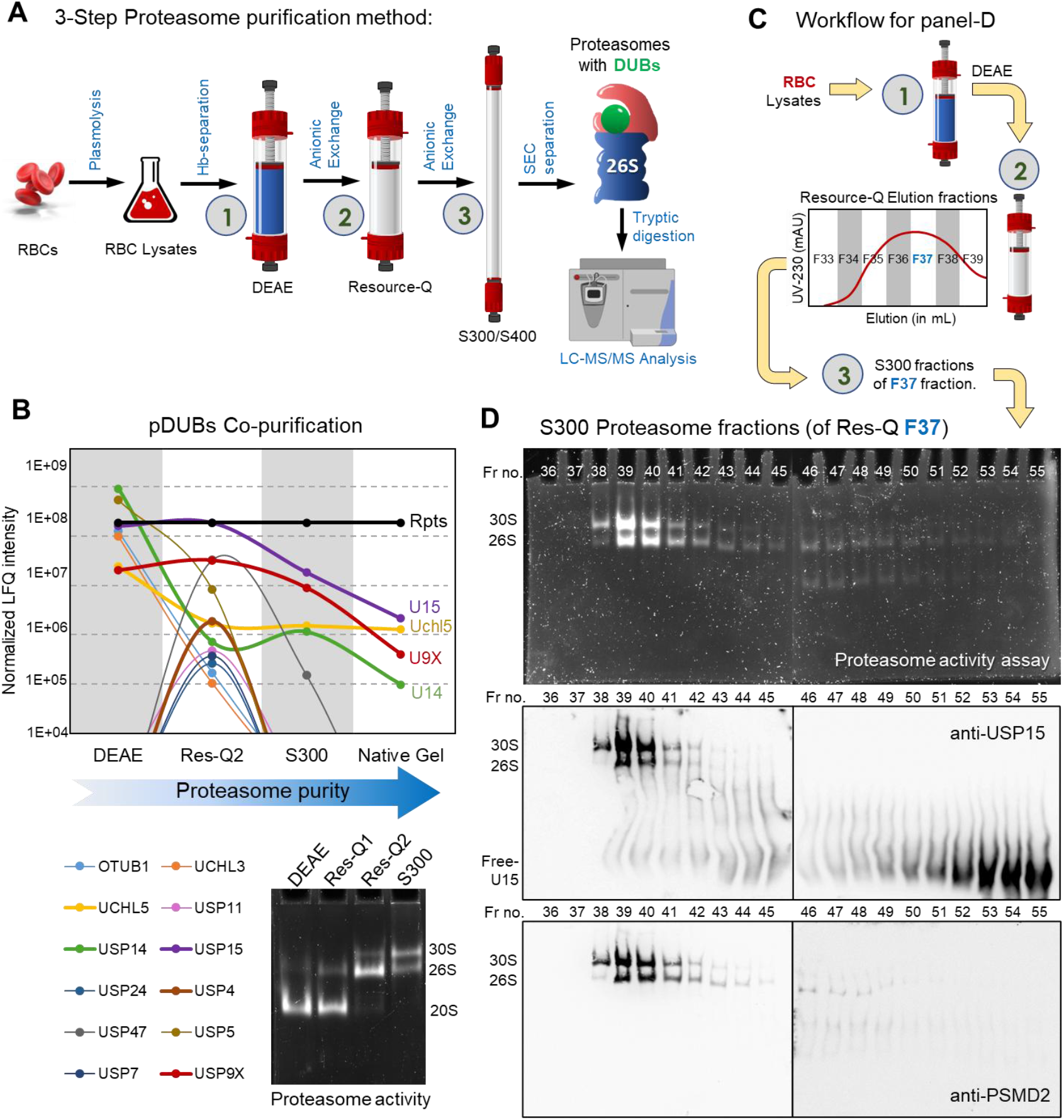
USP15 found to be associated with purified human 26S proteasomes. (A) Schematic representation of proteasome purification steps from human erythrocytes. The peak of proteasome peptidase activity from each step was pooled to the next step, and in parallel, the proteasome composition was analyzed by mass spectrometry (LC-MS/MS). (B) A graph displaying the total LFQ-intensity of DUBs found in each fraction, normalized to the LFQ-intensity average of all proteasome ATPase subunits (RPTs) found in the same fraction. Bottom right: The proteasome peptidase activity peak from each step was resolved by non-denaturing (“native”) gel, visualized by LLVY-AMC cleavage, demonstrating the proteasome configuration in each fraction. (C, D) All fractions eluting from the size exclusion column in step 3 were analyzed in parallel by Native Gel, visualized for peptidase activity (top), transferred to a PVDF membrane and stained for the presence of USP15 (middle) or proteasome subunit (bottom) to correlate between migration of USP15 and the proteasome complex.

To better understand the relationship of USP15 with the proteasome, we fractionated the resource Q elution to multiple proteasome-containing fractions and continued the purification protocol with each one separately (**Fig. 1C**). One of these fractions (F37 from panel C) confirmed a tight association between USP15 and 26S/30S proteasome when further resolved by size exclusion chromatography (**Fig. 1D**). USP15 distributed between two major pools: a high MW proteasome-bound, and a lower MW free form (**Fig. 1D, middle panel**). In summary, a systematic and comprehensive biochemical approach was instrumental in identifying USP15 as a genuine proteasome-associated DUB.

Having observed multiple DUBs in purified proteasome preparations led us to examine whether they cohabit on a single complex, or whether the sample is a heterogenous pool of distinct proteasome populations. Each proteasome-containing fraction after the Resource-Q step (**Fig. 1C; Sup Fig. 1)** was subjected separately to gel filtration fractionation. Comparing these parallel purified 26S proteasomes confirmed that it is possible to sort them according to their pDUB content (**Fig. 2A**). Whereas some proteasome fractions contained USP14 but not USP15, others contained USP15 but not USP14. Another fraction of proteasomes contained neither, indicating that these pDUBs are sub-stoichiometric to the overall proteasome population. In their respective fractions, a majority of USP14 or USP15 co-migrated with active 26S proteasome complexes by native gel (**Fig. 2B**). Subjecting the final proteasome fraction to MS/MS, confirmed that USP15 is by far the major pDUB in 26S^U15^ (with only trace amounts of USP14 or UCHL5), likewise, USP14 is the primary DUB in 26S^U14^ (**Fig. 2C**). By classifying 26S proteasomes according to the associated pDUBs, we have essentially identified a new subspecies of proteasomes containing USP15.

**Figure 2:**
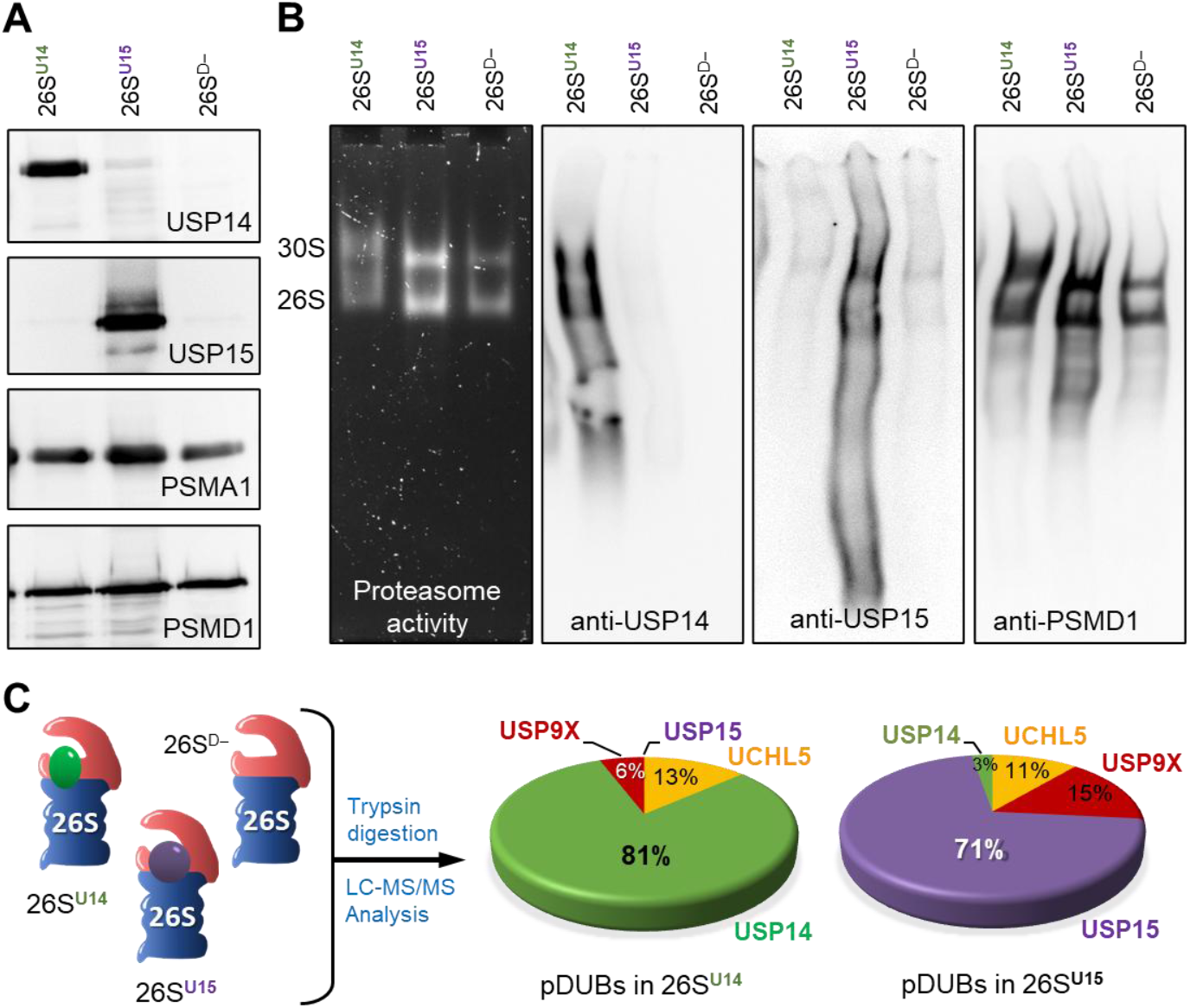
26S proteasome subspecies classified according to their pDUBs. Following the procedure from Figure 1C, each fraction eluting from the Resource Q column (step 2) was further subjected to a size exclusion column (step 3). In this manner, three proteasome pools were obtained, containing either USP14 (26S^U14^), USP15 (26S^U15^), or neither (26S^D-^). (A) SDS-PAGE immunoblotting was used to demonstrate the presence of USP14 or USP15 in each of the three proteasome pools. (B) Native gel activity (left) and Native Gel immunoblotting (middle, right) of the same samples demonstrating the co-migration of USP14 or USP15 with 26S and 30S proteasome species. (C) A pie chart displaying the distribution of the most abundant proteasome-associated DUBS (pDUBs) estimated by total LFQ-intensity in 26S^U14^ and in 26S^U15^ samples.

### USP15 associates with 26S proteasomes in mammalian cells

Having established the association of USP15 with proteasomes in red blood cells, we asked whether this represents a unique case or a broader phenomenon across cell types. We utilized differential centrifugation to rapidly separate proteasomes from whole cell extract (WCE), taking advantage of their large MW (~1.5-2.5 MDa for 26S and 30S complexes, respectively). A 100K RCF spin sedimented large particles such as ribosomes, followed by a 150K RCF to concentrate proteasomes (**Fig. 3A**) (Kuo et al., 2018). Both USP14 and USP15 cofractionated with the 26S proteasome in HEK293T and HeLa cells (**Fig. 3B**). Earlier findings reported that USP14 and its cognate Ubp6 in yeast can be found in both proteasome-bound and unbound pools (Elsasser et al., 2002; Guterman & Glickman, 2004; Hanna et al., 2006; Kuo & Goldberg, 2017; Leggett et al., 2002; Matiuhin et al., 2008; Rosenzweig et al., 2012). By the current method, USP14 also distributed in both high- and low-density fractions, yet in comparison, USP15 concentrated mostly with the proteasomes (**Fig. 3B**).

**Figure 3:**
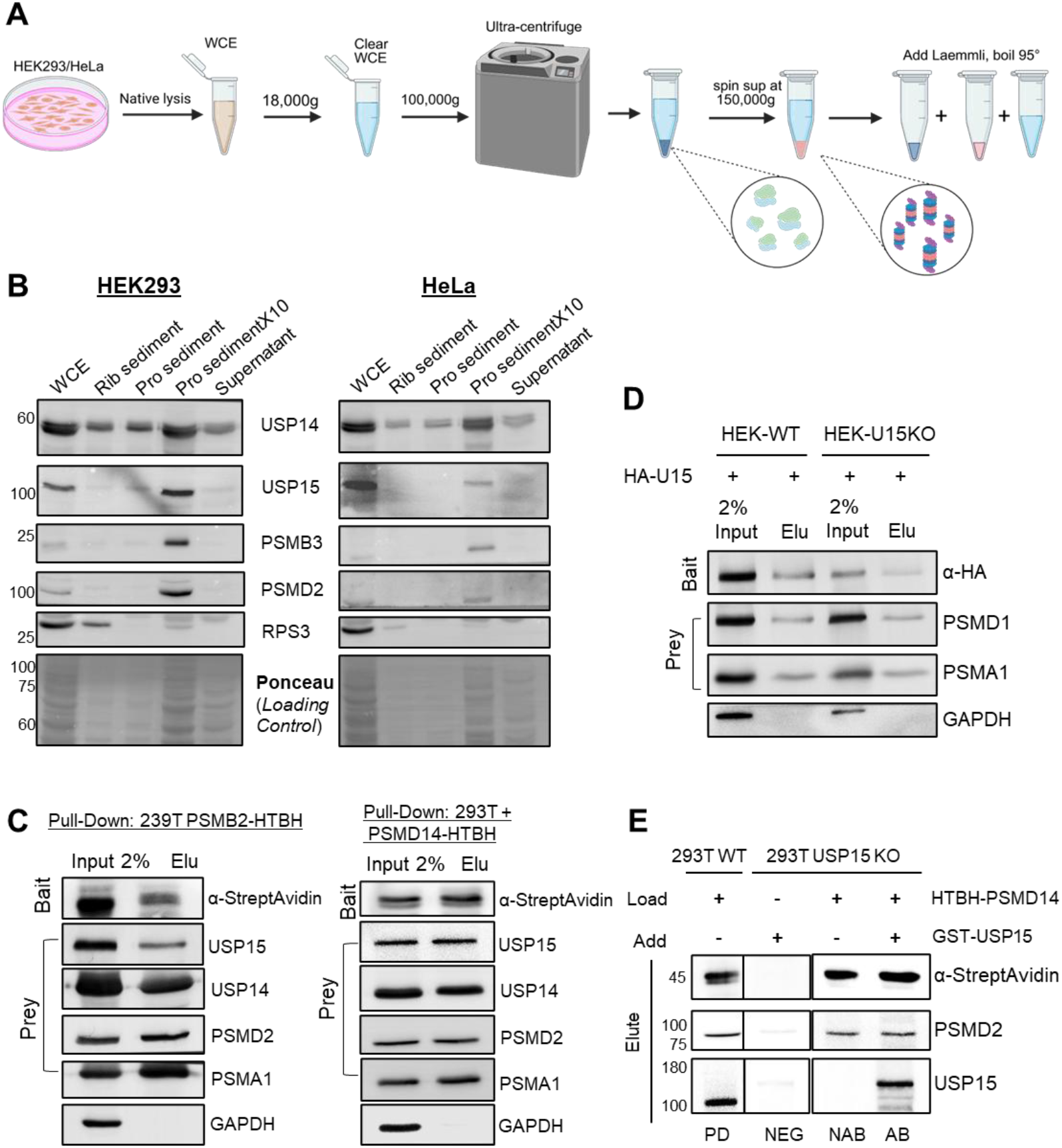
Interaction of USP15 with 26S proteasome in mammalian cells. (A) Schematic illustration of ultra-centrifuge spin protocol for proteasome enrichment. Created with BioRender.com. (B) SDS PAGE immunoblots of fractions of the ultra-centrifugation spin protocol obtained from HEK293T and HeLa cells. Pellet Rb-ribosomal represents enrichment fraction after 100K spin for 1h. Pellet Pr. Represent the proteasome-enriched fraction after a 150K spin for 1.5h. Pellet Pr(10X) is a 10X concentrated sample. Supernatant is the residual soluble material after all spins. USP15 and USP14 are detected in the proteasomal fraction together with 19S and 20S subunits. (C) Proteasomes from HEK293 cells endogenously expressing a biotinylated HTBH-tagged 20S subunit (PSMB2) or a transiently expressing HTBH-tagged 19S subunit (PSMD14) were pulled using Streptavidin beads to identify co-associating proteins. 2% input and elution fractions were subjected to immunoblot for the detection of USP15, USP14, a 19S subunit, a 20S subunit, or GAPDH as a non-proteasomal control. (D) Co-Immunoprecipitation of HA-tagged USP15. HA-USP15 was expressed in HEK293T or HEK293T USP15 KO cells, and immunoprecipitated by magnetic beads bound to HA antibody. 26S proteasomes were detected in the precipitated fractions using antibodies against a 19S subunit, a 20S subunit, or GAPDH as a non-proteasomal control. (E) USP15 addback to 26S proteasomes. 26S proteasomes from USP15 KO cells transiently expressing HTBH-tagged PSMD14 were immobilized on Streptavidin beads. Recombinant GST-USP15 was introduced, washed, and SDS-PAGE resolved eluted proteasomes for immunoblotting as indicated (right lane). Samples without addback confirm the lack of USP15. A similar procedure from USP15 KO cells, which do not express the tagged proteasome, demonstrates that USP15 does not spuriously bind to the column (second to left). As an indication of naturally associating USP15, proteasomes were similarly isolated from WT cells (left).

The presence of USP15 in high MW fractions of clarified cultured cell extracts suggests that it may interact with the proteasome. Direct association was confirmed by affinity co-purification. Initially, proteasomes were isolated directly from WCE either through His6-Tev-Biotin-His6 (HTBH)-tagged 19S (PSMD14) or 20S (PSMB2) particles, respectively (Choi et al., 2023). In either case, USP14 and USP15 coeluted with these affinity-purified proteasomes (**Fig. 3C**). Also, in a reciprocal approach, HA-USP15 co-immunoprecipitated both 19S and 20S subunits (**Fig. 3D**). Initially, we expressed HA-USP15 in CRISPR-KO cells to diminish potential interference of un-tagged endogenous USP15 for association with proteasomes yet later achieved similar results from WT cells (**Fig. 3D**). Interestingly, in either case, the USP15-associated pool of proteasomes also contained USP14, indicating that multiple pDUBs cohabit on some proteasomes simultaneously (**Fig. 3D**).

In order to validate whether the USP15-proteasome interaction is direct, we designed an assay to add-back recombinant USP15 to isolated proteasome complexes. 26S proteasomes lacking USP15 (from a KO cell line) were immobilized on Streptavidin beads, incubated with recombinant GST-USP15, washed, and eluted. USP15 co-eluted with these proteasomes but was not detected similarly treated chromatography beads without the presence of proteasomes (**Fig. 3E**). We concluded that exogenous USP15 can associate directly with purified proteasome complexes.

### The nature of the USP15-Proteasome association

Having established evidence that both endogenous and recombinant USP15 bind to the proteasome, we further investigated this association. USP15 was previously reported to bind and process K63, K48, and K11 Ub chains (Cornelissen et al., 2014; Han et al., 2024; Lange et al., 2024). Considering this information, it was unclear whether the association of USP15 can regulate the Ub levels on the proteasome. Conditions that elevate ubiquitinated substrates would help address this question. As expected, treating the cell culture with the proteasome inhibitor MG132 increased polyUB conjugates. This pervasive ubiquitination pattern was mildly intensified in USP15 KO cells relative to USP14 KO cells (**Fig. 4A)**. High-molecular-weight polyUb species were found to accumulate to a greater extent on proteasomes purified from USP15 KO cells than on those from wild-type or USP14 KO cells. (**Fig. 4B**). This supports the importance of USP15 as a proteasome-associated deubiquitinase (pDUB).

**Figure 4.**
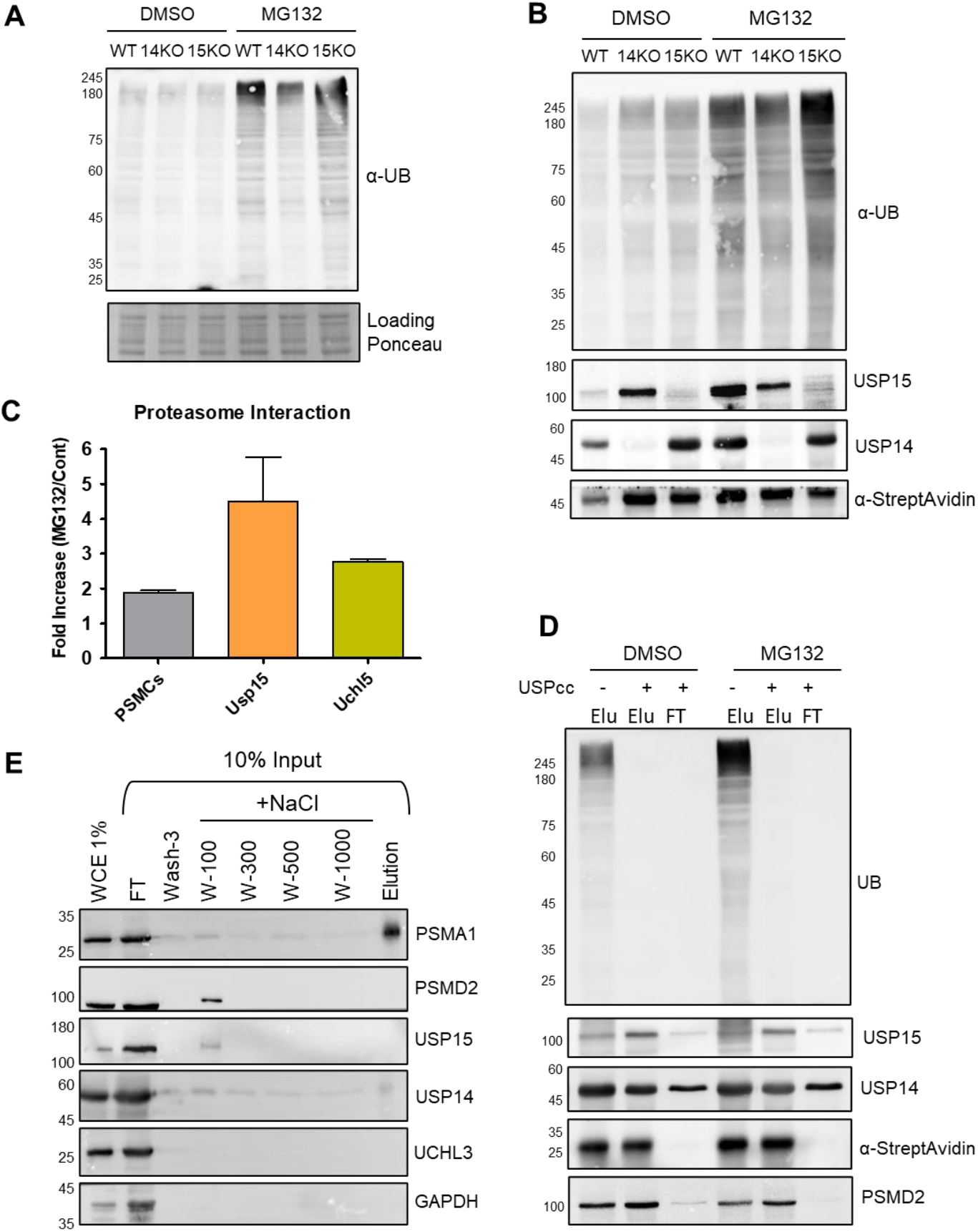
The relationship between the 19S-localized USP15 and polyubiquitin. (A) WCE from WT, USP14 KO or USP15 KO treated with either DMSO or 10 µM MG132 for 4 hours, were resolved by gradient (4%-20%) SDS-PAGE (GenScript M00652) and immunoblotted for total polyUb. Ponceau staining of protein content represents loading control. (B) proteasomes were isolated from HTBH-PSMD14 transfected WT, USP14 KO or USP15 KO treated with either DMSO or 10 µM MG132 for 4 hours and resolved by gradient (4%-20%) SDS-PAGE and immunoblotted for associated polyUb, USP15 and USP14. Streptavidin serves as a loading control for proteasome content. (C) Affinity-purified WT proteasome, as in (B), was subject to MS/MS. The ratio of USP15 content in proteasome from MG132-treated cells to untreated cells is compared to the similar ratio of UCHL5 and the average of all 6 RPT ATPases (PSMC). (D) To evaluate whether the association of USP15 is Ub-mediated, HTBH-PSMB2-immobilized proteasomes were treated with or without 1 µM USP2cc (catalytic core), a promiscuous DUB. Following treatment, the unbound fraction (FT) was separated, and the bound fraction was eluted with 1x SDS buffer. The various samples were stained for Ub, USP15, USP14, or PSMD2 to evaluate retention with the proteasome. Streptavidin serves as a loading control for the anchored 20S proteasome. (E) Anchored HTBH-PSMB2 proteasome was sequentially washed with increasing NaCl concentration in PBS buffer lacking ATP to disassemble 19S from 20S. The wash fractions were stained for PSMD2, USP15, or USP14 to evaluate retention with the proteasome. PSMA6 serves as a loading control for the anchored 20S proteasome. UCHL3 and GAPDH reflect proteins that are unrelated to proteasome.

Immunoblot analysis of MG132-treated proteasomes demonstrated that they retain their associated USP14 and USP15 (**Fig. 4B)**. The relative levels of USP15 increased by over four-fold in MG132-treated HTBH-PSMB2 affinity-purified proteasomes as calculated from MS/MS analysis (**Fig. 4C**). This observation cannot be explained solely by the assembly of additional 26S holocomplexes (as reflected by other 19S subunits; **Fig. 4C**). The abundance of polyUb conjugates with these MG132-treated proteasomes opens the possibility that they may play a role in recruiting USP15. To address this hypothesis, HTBH-PSMB2 proteasomes were affinity-purified, immobilised, and treated with USP2cc, a potent DUB with promiscuous specificity (Komander et al., 2009; Ryu et al., 2006), to remove associated Ub. Following this treatment, no trace of Ub was detected in eluted proteasomes, while retaining their associated pDUBS: USP14 and USP15 (**Fig. 4D**). In this respect, USP15 showed less sensitivity to Ub shaving than USP14 as some of USP14 did dissociate from proteasomes in the washes (**Fig. 4D; right lane**).

Tight association of USP15, which is not mediated through potential substrates in the form of Ub-conjugates, strengthens the conclusion that it binds directly to the proteasome. To pinpoint the subcomplex to which USP15 binds, we triggered disassembly of 19S RP from immobilized 20S CP (via HTBH-PSMB2) by depleting ATP and increasing salt concentration in the buffer (Leggett et al., 2002; C.-W. Liu et al., 2006). Low salt concentrations were sufficient to detach 19S subunits from the immobilized 20S subcomplex, and along with it, USP14 and USP15 (**Fig. 4E**). All these observations converge to support a direct association of USP15 to the 19S RP.

### A UBL domain in USP15 plays a role in proteasome interaction

The next objective was to provide some insight into the features of USP15 required for proteasome interaction. At this time, there is no high-resolution structure for full-length (FL) USP15. Crystal structures of partial USP15 constructs have been published (Harper et al., 2011; Teyra et al., 2019; Ward et al., 2018). Using an alpha fold model (AF-Q9Y4E8-F1), we assigned different structural domains onto the predicted tertiary structure (**Fig. 5A**). To determine which of the structural domains contribute to proteasome association, we cloned and purified the following fragments (**Sup. Fig. 2**): Full-length USP15, UBL1, the catalytic USP domain, UBL2, and a chimeric USP domain lacking the internal UBL2 (referred to herein as D1D2). Since the UBL of USP14 plays an important role in anchoring USP14 to the proteasome, we paid particular attention to the two UBLs embedded in USP15.

**Figure 5:**
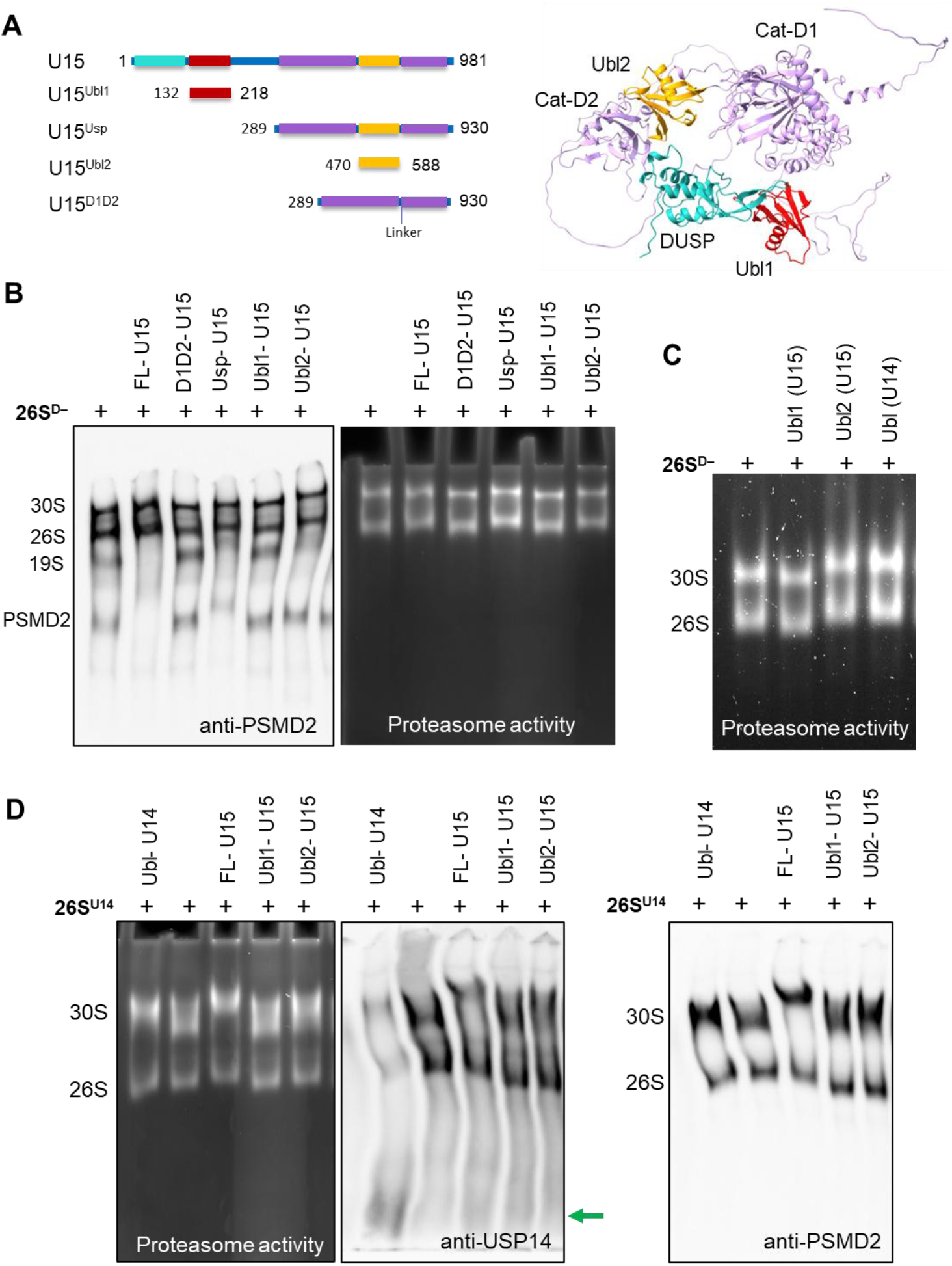
Domain mapping of the USP15 interaction with proteasomes. (A) Schematic illustration of putative USP15 structural domains, marked on the primary and AlphaFold3-predicted tertiary structures. Light blue: DUSP, red: UBL1, violet: USP (subdivided into D1 and D2), gold: UBL2. Full and truncated versions used in this study are listed within the appropriate amino acid borders. (B) Proteasome lacking transiently associated DUBs was incubated with USP15 constructs as indicated and resolved by 4% native gel. Proteasome proteolytic activity was visualized by cleavage of LLVY-AMC, and proteasome migration was evaluated by staining for 19S subunit PSMD2. Gel shift was noticeable by either detection method for 26S and 30S proteasomes incubated with FL-USP15, USP, or UBL2. A slightly more pronounced upward shift was visible for free PSMD2 and free 19S upon incubation with the same USP15 constructs. (C) Gel shift assay for proteasome with recombinant UBLs of USP15 or USP14. Gel electrophoresis was for a longer duration than in panel B in order to increase sensitivity to the gel shift with the small UBLs. (D) Competition assay for the proteasome bound to USP14. Proteasome containing USP14(26S^U14^) was incubated with UBLs of USP15 and USP14, or FL-USP15 as indicated, resolved by native gel and stained for either USP14 or PSMD2. Release of USP14 was detected only upon incubation with UBL of USP14. Upon incubation with FL-USP15, a noticeable gel shift was detected for proteasome retaining USP14, indicating the presence of both pDUBs simultaneously.

It is possible to detect the association of proteasomes with potential UBL-containing proteins by the migration pattern in non-denaturing (native) PAGE (Elsasser et al., 2002, 2004). Initially, 26S lacking pDUBs (26S^D-^) were incubated with recombinant USP15 and resolved by native PAGE. Upon addition of USP15, free 19S and other PSMD2-containing particles were undetected by native immunoblotting, likely reflecting a shift in their migration due to interaction with USP15 (**Fig. 5B**; lanes 1,2). An upward shift of these two species was noticeable upon the addition of the USP domain, in line with the larger MW of the resulting complexes; however, the D1D2 construct lacking the internal UBL2 domain did not exhibit the same behavior (**Fig. 5B**; lanes 3,4). Incubation with UBL2 led to a visible change in migration of the 19S, whereas addition of UBL1 domain showed no discernible effects (**Fig. 5B**; lanes 5,6). Upon longer resolution, a visible gel shift of the holocomplexes (26S and 30S) became apparent upon addition of UBL2 but not of UBL1 (**Fig. 5C**). This outcome mimics the effect of the USP14 UBL domain (**Fig. 5C**; lane 4), a known interactor of proteasome.

Next, we wished to test whether the UBL domains of USP14 and USP15 compete for the same site on proteasomes. USP14 UBL domain alone was sufficient to knock off the endogenously bound USP14 from the proteasome (**Fig. 5D**; lane 1). By this method, none of the USP15 domains outcompeted USP14 from 26S^U14^ proteasomes (**Fig. 5D**). In a complementary experiment, USP14 UBL was also found to be inefficient in replacing endogenous USP15 (**Sup. Fig. 3**). We conclude that although these two pDUBs contain UBL domains, the nature of their association with the proteasome is not equivalent.

### USP15 UBL domains do not compete with USP14 for the same interaction site on PSMD2

The primary proteasome UBL receptor is the large subunit, PSMD2. Structurally, PSMD2 can be roughly divided into three domains: an extended N-terminal α-helical domain, a circular solenoid-like PC domain, and a loosely folded C-terminal region (Shi et al., 2016). Previous studies have shown that the PC domain can bind Ub, PolyUb, and UBL domains from various UBL-containing proteins (Chen et al., 2016; Guo et al., 2011). Additionally, the N-terminal domain has also been reported to associate with diubiquitin (diUb) (Boughton et al., 2021). We note that the USP14-UBL has a designated binding site on PSMD2 (Shi et al., 2016). Since UBLs of both USP14 and USP15 were potentially able to associate with the proteasome simultaneously (**Fig. 5D**), we wished to explore the USP15-UBL binding site. We used AlphaFold3 (AF3) to model potential interactions by introducing the full-length amino acid sequence of PSMD2 alongside various UBL domain sequences. For each UBL 100 models were generated. To evaluate the credibility of the predicted models, we first modeled the interaction between PSMD2 and the UBL domain of USP14 and compared the AF3-predicted complex to the cryo-EM structure of the USP14-UBL-bound proteasome (PDB: 7W37). **Fig. 6A** depicts the top five models (of 100) of PSMD2 with USP14-UBL. Importantly, not only does AF3 predict de novo folding of the entire PSMD2 sequence accurately with respect to the experimental structure, it also positions USP14-UBL at the reported binding site with precise orientation (**Fig. 6B**). We then applied the same strategy to model the interaction between PSMD2 and the UBL domain of RAD23A, another well-established proteasome interactor. Notably, AF3 correctly positioned Rad23A-UBL at T1 (Shi et al., 2016) and interestingly also at the second known UBL binding site, T2 (**Fig. 6C,D**).

**Figure 6.**
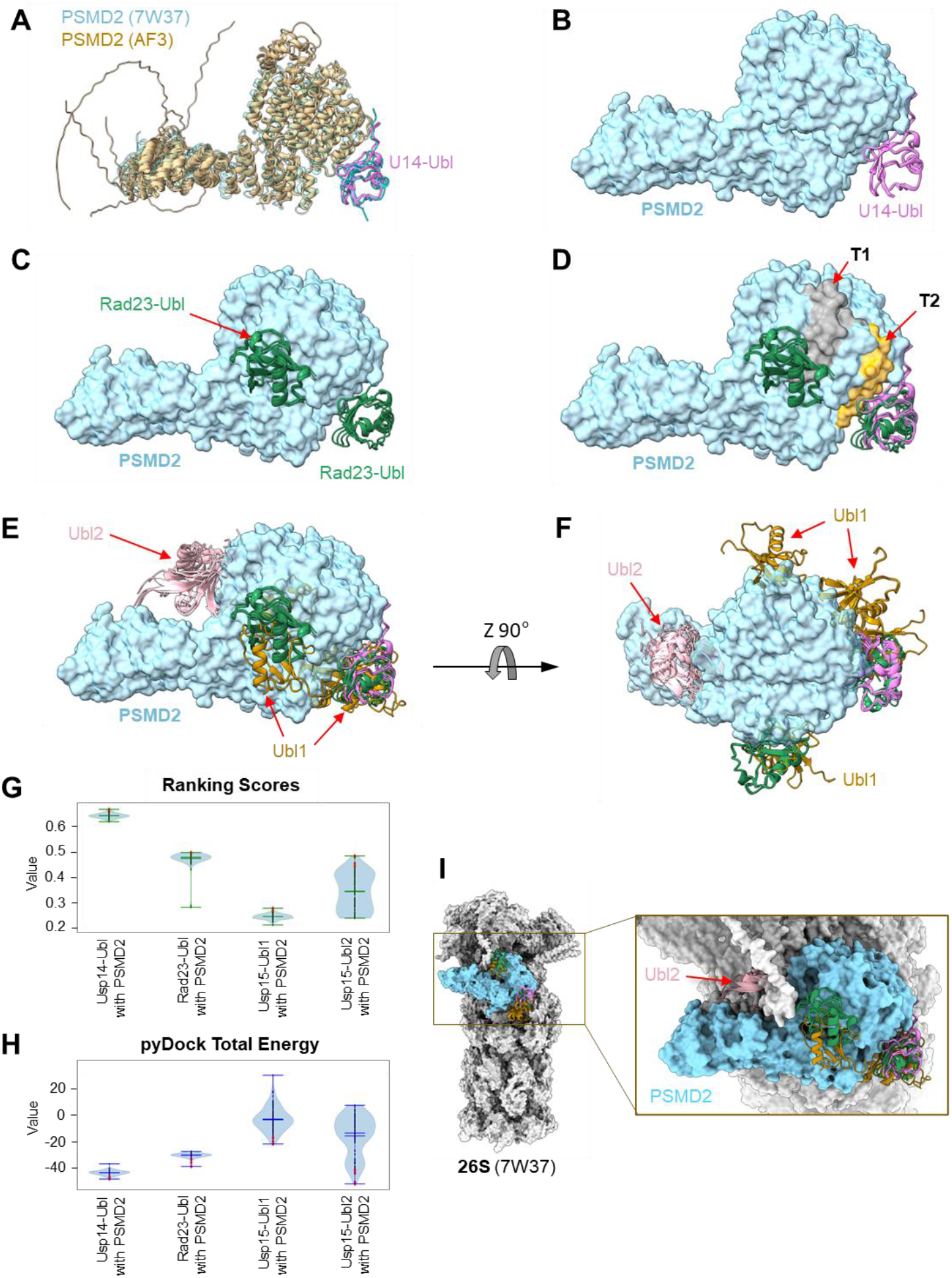
AlphaFold3 (AF3) models for PSMD2 with various UBL domains. (A) AF3 best five modeled complexes of PSMD2-UBL(U14) shown in tan and pink ribbon, respectively, superimposed on PSMD2-UBL(U14) from a cryo-EM structure (PDB 7w37; chain x in transparent cyan ribbon, chain f in cyan ribbon) shown for reference. (B-G) AF3 PSMD2-UBL five best ranking complexes superimposed on 7w37 PSMD2 (Note: PSMD2 from the AF3 models was removed for clarity). PSMD2 from 7w37 is shown in cyan surface while all predicted AF3 UBL models are shown in different colored ribbons. (B) AF3 PSMD2-UBL(U14) modeled complexes; the UBL in pink ribbon. (C) AF3 PSMD2-UBL(RAD23) modeled complexes; the UBL in green ribbon. (D) T1 and T2 regions on PSMD2 are colored in grey and orange, respectively. UBL(U14) (pink ribbon) and UBL(RAD23A) (green ribbon). (E) As in D, with the addition of UBL1(U15) (brown ribbon) and UBL2(U15) (light pink). (F) As in E, rotated x 90 and y −30 in ChimeraX for an additional perspective. (G) Violin plots showing AF3 model ranking scores for all 100 AF3 models of PSMD2-UBL complexes of USP14, RD23A, and USP15 (UBL1 and UBL2). Best five ranking scores are shown in red dots. (H) Same as in G, but presenting pyDock total energy respectively (I) As in F, in the context of the full proteasome (from 7w37), PSMD2 is colored in cyan, while all other proteins in PDB 7w37 are colored in grey. The insert shows a zoom in to PSMD2 area.

The promising predictions of AF3 encouraged us to apply the same workflow to assess potential interactions between the two USP15-UBL domains and PSMD2. Each UBL domain (UBL1 and UBL2) was modeled independently with PSMD2. AF3 modelling of UBL1 produced highly variable and inconsistent binding orientations across the predicted structures (**Fig. 6E,F**). By contrast, UBL2 produced more consistent models, clustering at a single binding site, distinct from the T1 and T2 sites (**Fig. 6E,F**). The top five models of UBL2 exhibited a high-ranking score with low binding energy, as computed by PyDock, similar to USP14-UBL and Rad23A-UBL (**Fig. 6G,H**). Nevertheless, when considering all 100 generated models, the variability was substantial, significantly higher than that of the USP14 or Rad23A UBLs. When considered the predicted binding site within the framework of the entire 26S holocomplex, the orientations of USP15-UBL2 appeared sterically unfavorable (**Fig. 6I**). Specifically, upon incorporation of PSMD2 into the 19S subcomplex, the newly predicted binding sites for USP15 UBLs on PSMD2 would be buried within the proteasome, largely inaccessible to the entire USP15 protein. Therefore, AF3 predictions suggested limited interaction potential between USP15 and PSMD2.

### USP15 contributes to substrate selection by disassembling short polyubiquitin chains on the proteasome

Having established that USP15 is a proteasome-associated DUB, we then turned to investigate the mechanistic contribution of USP15 on the fate of polyubiquitinated substrates. USP15-enriched proteasomes, 26S^U15^, were incubated with a published polyUb-conjugate with a fixed non-hydrolyzable proximal bond that is resistant to the Rpn11 DUB activity (**Sup. Fig. 4**) (Singh et al., 2016). 26S^U15^ trimmed di- or tri-Ub-Globin from their distal end, releasing shorter intermediates (**Fig. 7A**). Treatment with a pan-cysteine modifier such as iodoacetamide (IAA) blocked Ub trimming (**Fig. 7A;** lanes 4-6), confirming that disassembly of di- or tri-Ub in this setup is attributed almost exclusively to cysteine-based pDUBs such as USP15. Therefore, proteasomes lacking USP15 (26S^D-^) displayed no detectable trimming of these substrates (**Fig. 7A;** lanes 9,10). In agreement, treating 26S^U15^ with a Zn^2+^-chelator (1,10-ortho-phenanthroline; O-PHEN) to inhibit Rpn11/PSMD14 did not influence processing of these conjugates (**Fig. 7A;** lane 7).

**Figure 7.**
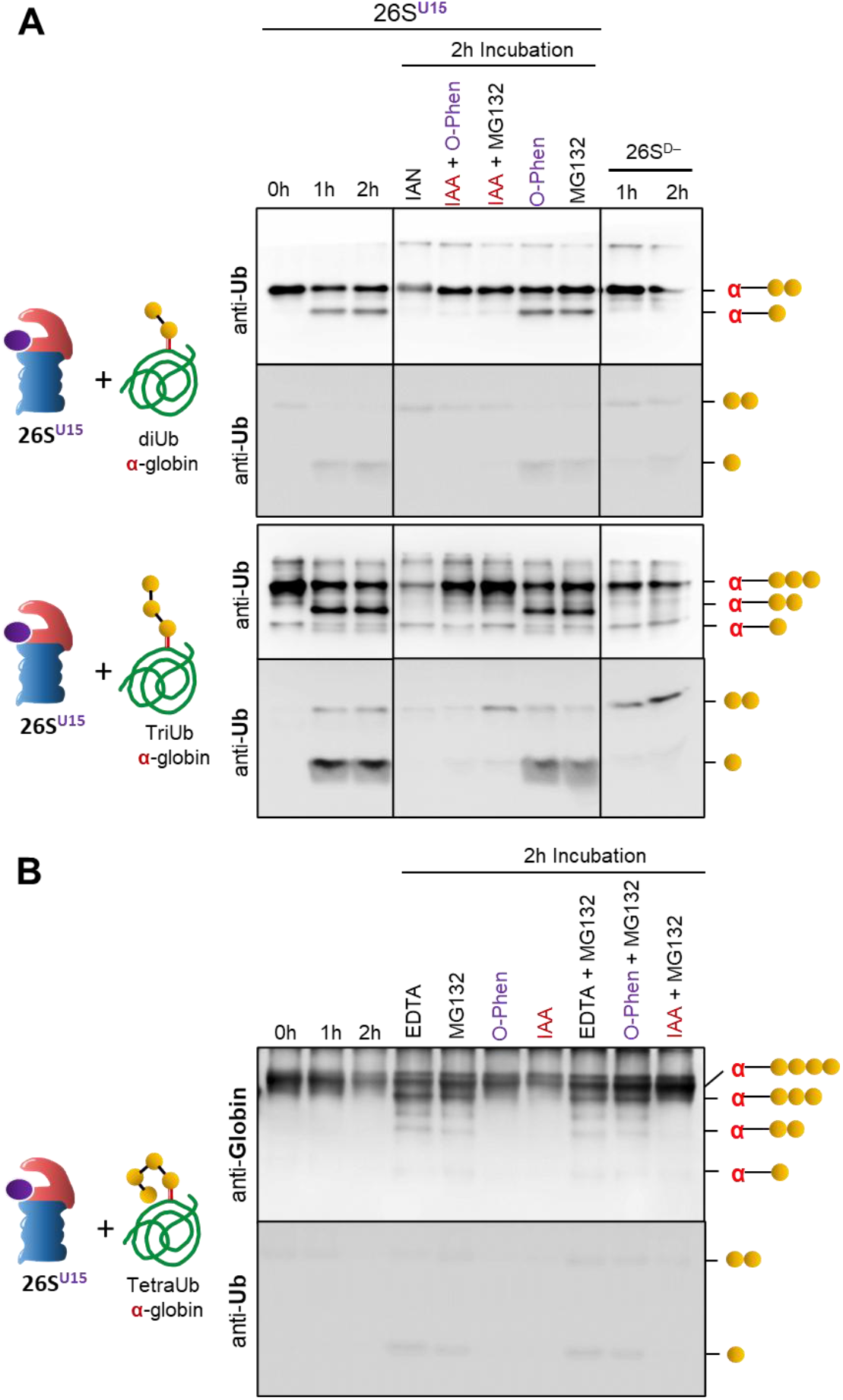
USP15-bound proteasomes process polyubiquitinated globin. (A) Purified human 26S proteasomes containing USP15 (26S^U15^) or 26S proteasomes without USP15 (26S^D-^) were incubated with diUb-α-globin or triUb-α-globin substrates at a 1:100 molar ratio for up to 2 hours with or without inhibitors as labeled. IAA, Iodoacetamide; O-PHEN, 1,10-Ortho-Phenanthrolin, MG132, EDTA. Reactions were terminated and products resolved by Tris-Glycine 15% SDS-PAGE and stained for Ub. Reaction components are marked on the right with a red α for Globin, and a gold dot for each Ub molecule. (B) Same as A but 26S^U15^ or 26S^D-^ were incubated with tetraUb-α-globin at a 1:100 molar ratio for up to 2 hours with or without inhibitors as labeled. Reactions were terminated and contents resolved by Tris-Glycine 15% SDS-PAGE and immunoblotted with either anti-globin (upper panel) or anti-Ub (lower panel).

In contrast to di- and tri-Ub-Globin, no trimming was observed for tetra-Ub-Globin by 26S^U15^ within the same time frame (**Fig. 7B**, lanes 1-3). This result indicates that the K48-linked tetraUb modification is relatively resilient to USP15-associated proteasomes, leading to degradation of the conjugated substrate. Since di- and tri-Ub-Globin were trimmed by USP15-bound proteasomes, these conjugates were largely rescued from degradation by 26S^U15^. By contrast, 26S^D-^ partially degraded the same substrates, along with the covalently attached Ub (**Fig. 7A**; lane 10, **7B**, lanes 1-3). We propose that the degradation of Ub-Globin by 26S^D-^ is due to the absence of trimming activity, as inhibition of USP15 in 26S^U15^ similarly led to degradation of these substrates (**Fig. 7A;** lane 4). Proteolysis was confirmed to be proteasome-dependent, as it was inhibited by MG132 or EDTA (**Fig. 7A**, lane 6; **Fig. 7B**, lanes 4,5,8,9,10). Chain trimming was uncoupled from proteolysis, indicating it is an upstream, preparatory event. We conclude that trimming of di- or tri-Ub was quicker than proteolysis of the conjugated Globin by 26S^U15^ in this setup; hence, USP15 can rescue substrates from degradation in contrast to tetraUb-Globin for which substrate degradation by 26S^U15^ is faster than chain processing.

Next, we tested the behaviour of 26S^U15^ on a set of substrates in which all Ub linkages are isopeptide bonds: Ub-cyclin B1, diUb-Cyclin B1, and tetraUb-Cyclin B1 (**Fig. 8**). The primary outcome of proteasome action on Ub-Cyclin B1 was deubiquitination and release of Cyclin B1 (**Fig. 8A**, lanes 1-3). By differentially inhibiting USP15 (IAA) or Rpn11 (O-PHEN), we concluded that both USP15 and Rpn11 contributed to this outcome (**Fig. 8A**, lanes 4-6). Similarly, for diUb-Cyclin B1, shaving of the diUb chain can be attributed to Rpn11, whereas trimming of a single Ub unit from the distal end was attributed to USP15 (**Fig. 8B**, lanes 1-6). Proteasome lacking USP15, 26S^D-^, exhibited no trimming activity on these substrates, but interestingly, “shaved” diUb was not accompanied by free Cyclin B1 (**Fig. 8B**, lower panel lanes 7,8). This result suggests that Cyclin B1 was degraded by 26S^D-^. Similar degradation was observed for diUb-Cyclin B1 by 26S^U15^ upon inhibition of USP15 (**Fig. 8B**, lower panel lane 4).

**Figure 8.**
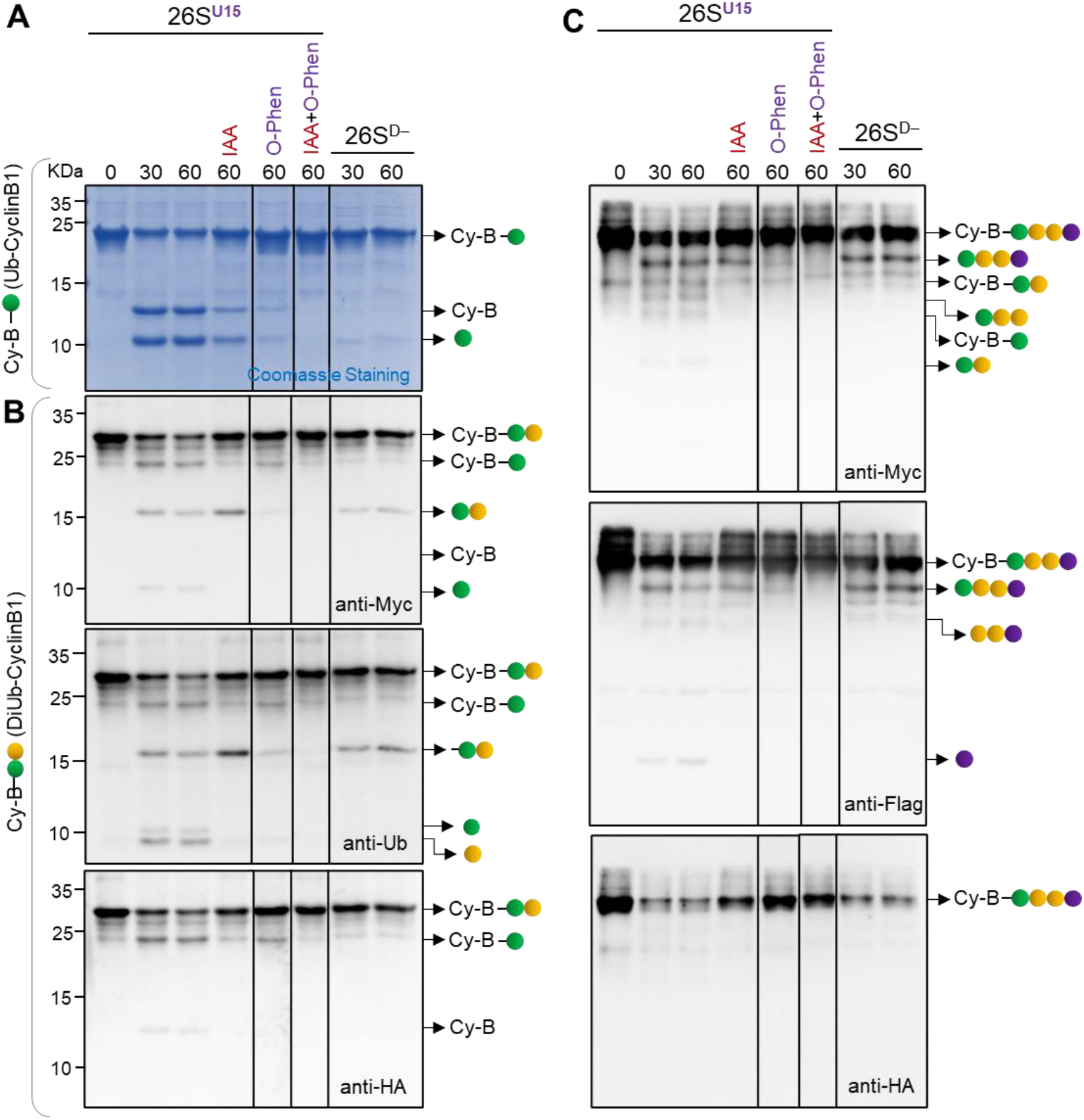
USP15-bound proteasomes process a panel of polyubiquitinated cyclin B1. Purified human 26S proteasomes containing USP15 (26S^U15^) or 26S proteasomes without USP15 (26S^D-^) were incubated with a panel of differentially ubiquitinated Cyclin B1 (monoUb-, diUb- or tetraUb-cyclin B1) at 1:100 molar ratio for up to 1 hour. In all cases, Cyclin B1 was N-terminally HA-tagged, and the proximal Ub was N-terminally Myc-tagged (green). The distal Ub unit of the tetraUb modification was N-terminally Flag-tagged (purple). Other Ub units are illustrated as gold dots. Inhibitors were added to some of the reactions as indicated: IAA, Iodoacetamide; O-PHEN, 1,10-Ortho-Phenanthrolin. Reactions were terminated and products resolved by Tris-Glycine 15% SDS-PAGE. (A) Coomassie staining of reaction with monoUb-Cyclin B1. (B) Reaction of diUb-Cyclin B1 immunoblotted with anti-Myc, anti-Ub or anti HA. (C) Reaction mix of tetraUb-Cyclin B1 substrates immunoblotted with anti-Myc, anti-Flag or anti HA.

Incubation of K48-linked tetraUb-Cyclin B1 with 26S proteasomes resulted in degradation of Cyclin B1 with simultaneous release of the conjugated K48-linked tetraUb irrespective of the presence or activity of USP15 (**Fig. 8C**, lanes 1,2,3,7,8). Very little trimming was observed, confirming once again the resilience of tetraUb to processing by the proteasome. The release of tetraUb was attributed to Rpn11 shaving activity (**Fig. 8C**, lanes 4-6). For this K48-linked tetraUb-conjugate, the data suggests that shaving occurred after the substrate commitment to degradation, as we did not detect liberated substrate (which was observed with shorter chains). To summarize, 26S^U15^ differentiates between K48-linked tetraUb and shorter Ub modifications, decreasing the commitment to degradation of shorter conjugates.

## Discussion

In this study, we expand the repertoire of DUBs associated with proteasomes into a family of pDUBs. USP15 joins as a new member, alongside UCHL5 and USP14. Each pDUB uniquely contributes to proteasome function. Proteasome-associated USP15 enables differentiation between substrates conjugated with short vs long chains (**Fig. 9**). Specifically, USP15 rapidly processed chains shorter than tetra-Ub, filtering them out of the downstream proteolytic path. The resistance of K48-linked tetraUb to USP15 DUB activity committed the conjugates to degradation. Proteasome-associated USP14 has been demonstrated to remove supernumerary K48-linked Ub chains (Lee et al., 2016), and UCHL5 has been ascribed to prune branched K48-containing chains (Deol et al., 2020). The three pDUBs appear to orchestrate preparatory steps depending on the Ub topology on the substrate. This fits their peripheral position on the 19S, contributing to their enzymatic activities as required. Rpn11, adjacent to the substrate translocation channel, executes the removal of the residual Ub once the substrate is committed to degradation (Verma et al., 2002; Worden et al., 2017). Although they undoubtedly share overlapping functions, they are not equivalent, and each adds exclusive properties to the proteasome, adding layers of regulation to substrate selection.

**Figure 9.**
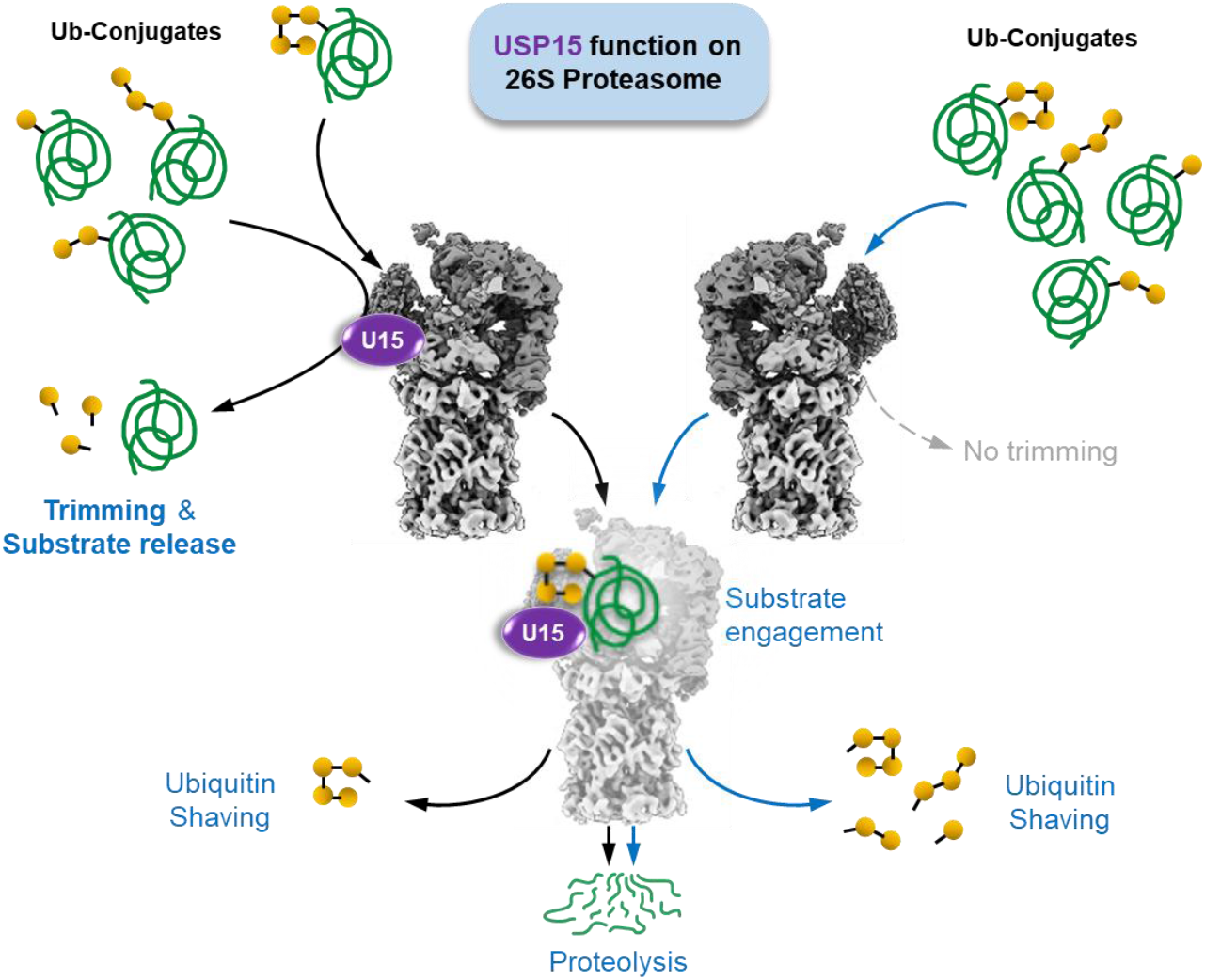
USP15 on 26S proteasomes selects polyubiquitinated conjugates for further processing. Model figure shows the role of USP15 on the 26S proteasomes to differentially process shorter Ub-chain conjugates by trimming and TetraUb-conjugates by degradation. The 26S proteasomes devoid of any Cys-based pDUBs can proteolyse all Ub-conjugates shown but at different rates.

To fulfill their respective roles as transient interactors of the proteasome, each pDUB should have a defined docking site to position itself in an appropriate orientation. Since we confirmed that USP15 did not require Ub conjugates for proteasome association, it implies an intrinsic affinity. In the case of USP14/Ubp6, the UBL domain participates in anchoring to a defined Ub/UBL-binding site on PSMD2/Rpn1, though CryoEM highlights additional contact sites beyond the UBL (Dash et al., 2024; Huang et al., 2016). UCHL5 has no UBL domain, yet it too interacts at a specific location on the proteasome (Deol et al., 2020; Sakata et al., 2012). Our findings suggest that USP15 binds to the 19S, although it does not compete for the same binding site as USP14. Domain mapping of USP15 showed that UBL2 is an important structural element for binding the proteasome; in fact, AF3 predicted interaction on PSMD2, though not at the established conventional Ub/UBL binding sites. The steric hindrance at the predicted site questions the precise orientation. Previous reports suggest USP15 interacts with the COP9-signalosome (Dubiel et al., 2020; Meister et al., 2019), which lacks a documented Ub/UBL binding subunit. Accordingly, we propose that other domains of USP15 and other proteasome subunits probably contribute to proper positioning.

Proteasome-interacting proteins (PIPs) are generally transient and sub-stoichiometric. These include substrate shuttles (Zientara-Rytter & Subramani, 2019), stabilizers (Choi et al., 2023), phosphatases (Guo et al., 2011), E3 ligases (Buel et al., 2020), and DUBs such as USP15. As part of a broad dynamic sphere of interactions revolving around the 26S proteasome, additional DUBs may be awaiting reclassification as pDUBs. The game of “musical chairs” between PIPs enables adaptation of the proteasome to a variety of needs in different cell types. Erythrocytes appear to be abundant in UPS components, in particular USP15 and USP15-associated proteasomes. The discrimination of 26S^U15^ against shorter chains may help protect “poorly ubiquitinated” substrates against unwarranted protein degradation in these non-nucleated cells. In dividing cells or in cancer cells, excessive activity of USP15 may come with the cost of stabilizing oncoproteins and act as a pro-oncogenic factor (Xu et al., 2018; Zhong et al., 2021). In contrast, USP15 can also act as a stabilizer of tumor suppressors like P53 and IκBα (W.-T. Liu et al., 2017; Zheng et al., 2019). In human dopaminergic neurons, USP15 has been shown to enhance mitophagy by blocking Parkin-dependent mitochondrial ubiquitination (Cornelissen et al., 2014). While in Parkinson’s disease, USP15 cellular localization regulates autophagy through AKT-mediated phosphorylation, contributing to the progression of the disease (Wu et al., 2025). The new information that a portion of USP15 is localized to the proteasome elicits a need to reinterpret earlier observations linking it to cancer progression, mitophagy regulation, and autophagy in Parkinson’s disease, potentially placing these observations within a proteasome-dependent context.

## METHODS

### Cell culture

HEK293T cells were purchased from ATCC (Manassas, VA) and maintained in Dulbecco’s Modified Eagle Medium-DMEM (Sigma-Aldrich #D5796) supplemented with 10% FBS (Sigma-Aldrich #F7524) L-glutamine, sodium pyruvate, and penicillin/streptomycin (Sigma-Aldrich; G7513, # 105477, #P4333; 1:100).

### Cell transfection

For transfecting of HEK293T cells, 1:10 diluted polycation Polyethylenimine (PEI; Sigma-Aldrich #408727) reagent was mixed with the desired plasmid in ratio of 1:2, in order to optimize transfection efficiency cells were transfected in DMEM media without FBS. After 6 hours, the media was replaced with full DMEM media containing 10% FBS. The transfected cells were then incubated for 24 hours at 37°C for optimal protein expression.

### Cloning

DNA fragments of interest were amplified by polymerase chain reaction (PCR) using high-fidelity DNA polymerase (Phusion, NEB M0530) and gene-specific primers containing appropriate restriction sites or overhangs for ligation. PCR products were purified using a PCR cleanup kit (Zymo D4001) and digested with restriction enzymes (Thermo Scientific) when necessary. Purified inserts were ligated into linearized plasmid vectors using T4 DNA ligase (New England Biolabs, M0202) or by GIBSON assembly (EURx, E1050), according to the manufacturer’s protocols. Ligation mixtures were transformed into chemically competent E. coli (DH5α) by heat shock at 42°C for 1 min. Transformed bacteria were plated on LB agar supplemented with the appropriate antibiotic and incubated overnight at 37°C. Colonies were screened by colony PCR or restriction digest, and positive clones were verified by Sanger sequencing. All plasmid constructs were prepared using a miniprep kit (Promega, A1460) and quantified by spectrophotometry (Nanodrop). All DNA design was performed using Benchling [Biology Software]. (2025). Retrieved from https://benchling.com (RRID:SCR_013955).

### Western blot

Proteins were resolved on 8%, 10%, 15%, or 4-20% tris-glycine gels, 20 μg of WCE, and 25% of Elution in each experiment. Followed by transfer onto nitrocellulose (TAMAR #10401380) or 0.45µm polyvinylidene difluoride (PVDF; Sigma-Aldrich #IPVH00010). The membranes were then blocked in a 5% dry milk dissolved in TBS-T to prevent nonspecific binding of antibodies. Following blocking, the membranes were probed with specific primary antibodies targeting the proteins of interest, incubated 1 hour at room temperature. The membranes were washed with TBS-T to remove excess primary antibody and then, as needed, incubated with corresponding secondary antibodies (a-Ms, a-Rb) conjugated to HRP for enhanced detection. Protein bands were visualized using ECL and captured using an imaging system, Fusion enhanced chemiluminescence.

### Co-immunoprecipitation with HA beads

Co-IP assay was performed using HA magnetic beads (MCE, HY-K0201) HEK293T cells were transfected with 3xHA-USP15 (pCDNA3 vector). Briefly, cells were lysed with buffer A (25 mM TRIS 7.4 pH, 10% Glycerol, 10 mM MgCl^2^, 1 mM ATP, 1 mM DTT). Beads were washed with 500μl of buffer A. Washed beads were incubated with 800 μg of clear cell lysate for 2 hours at 4°C, allowing for the specific capture of HA-tagged protein. After incubation, the flow-through was removed, and the beads were extensively washed 3 times with 500 μL of buffer A to remove nonspecifically bound proteins and contaminants. Following the washing steps, the protein complexes bound to the HA beads were eluted using 1 hour incubation with 2 mg/ml HA-free peptide (GenScript, RP11735-1). Subsequently, the eluted samples were subjected to Western blotting to identify the interacting proteins.

### Pull-Down with Streptavidin magnetic beads

HEK293T HTBH-PSMB2 stable cells (Choi et al., 2023) were lysed as described in the previous method. The beads were pre-washed with buffer A. The washed beads were incubated with 800 µg of clear cell lysate at 4°C for 1 hour. This allows the biotinylated HTBH tagged proteins in the lysate to bind to Streptavidin magnetic beads (GenScript, L00936). After the incubation supernatant was discarded, followed by wash with buffer A to remove nonspecifically bound proteins. The proteins bound to the Streptavidin beads were eluted using 100 μl of 1x Laemmli buffer and boiled for 5 min at 95°C. Streptavidin antibody was used to detect IP Proteasomes eluted from the beads in western blot. Subsequently, the eluted samples were subjected to Western blotting to identify the interacting proteins.

### Native gel, in-gel activity assay, and native immunoblotting

Native PAGE was performed to analyze the integrity and composition of proteasome complexes under non-denaturing conditions. Proteasome-containing samples were prepared in native loading buffer (0.45 mM TRIS base, 0.45 mM boric acid, 12.5 mM MgCl, 2.5 mM EDTA, Xylene cyanol). Samples were loaded onto 4% native polyacrylamide gels prepared in-house using acrylamide/bis-acrylamide (19:1) in native gel buffer (0.45 mM TRIS base, 0.45 mM boric acid, 12.5 mM MgCl, 2.5 mM EDTA. 1 mM ATP, 1 mM DTT). Electrophoresis was carried out at 4°C in ice cold running buffer (100 mM TRIS base, 100 mM boric acid, 2.5 mM MgCl, 1 mM EDTA. 0.5 mM ATP, 0.5 mM DTT) at a constant 120 V for ~3 hours. For In-gel activity assay, the gel was incubated in buffer A with 0.0125 mM Suc-LLVY-AMC (Bachem 4011369) at 37 °C for 10 min and imaged under UV light. For Native immunoblotting, the gel was soaked in SDS-running buffer for 20 min and then transferred onto 0.45 μm PVDF membrane using a wet transfer system.

### USP2cc Ub shaving

HEK293T expressing HTBH tagged proteasomes were treated with either DMSO or 10 μM MG132 for 4 hours. Cells were then lysed in lysis/wash buffer (25 mM TRIS 7.4 pH, 10% Glycerol, 10 mM MgCl2, 0.01% NP-40, 1 mM ATP, 1 mM DTT). 1 mg of clear lysate was added to pre-washed Streptavidin magnetic beads, and kept for 1.5 hours incubation at RT. Following 4x wash with wash buffer, 1 μM of recombinant HIS-USP2cc was added to the proteasome bound beads for 1 hour incubation at 37◦C. The un-bound fraction (flow through-FT) was kept, and the beads were extensively washed. Elution of the bound fraction was done by adding 60 μL of 1XSDS-loading buffer to each sample and boil for 5 minutes in 95◦C. From each fraction (FT and elution) 25% was loaded on SDS-gradient gel (4%-20%).

### CRISPR KO

Gene knockout was performed using the CRISPR/Cas9 system based on the S. *pyogenes* Cas9 nuclease. Guide RNA (gRNA) sequences targeting the gene of interest were designed to precede an NGG PAM motif and were cloned into the BbsI site of the Px459 expression vector (Addgene 62988), which encodes both Cas9 and a chimeric single-guide RNA (sgRNA). Oligonucleotides were phosphorylated, annealed, and ligated into the linearized plasmid backbone following the Zhang Lab protocol (Cong et al., *Science* 2013). Constructs were verified by Sanger sequencing and verified plasmids were transfected into HEK293T. After 48 hours, cells were either subjected to 5 μM puromycin (Sigma-Aldrich P8833) selection for enrichment for 7-12 days subsequently, clonal populations were established by limiting dilution, and knockout efficiency was validated by immunoblotting.

### Proteasome disassembly

HEK293T cells expressing HTBH-tagged proteasomes were lysed in PBS (Sigma-Aldrich D8537). A total of 2.5 mg of clarified lysate was incubated with pre-washed Streptavidin magnetic beads for 1 hour at room temperature with gentle mixing. The flow-through was collected, and the beads were washed three times with PBS. For salt washes, PBS containing NaCl at final concentrations of 100 mM, 300 mM, 500 mM, and 1 M was sequentially added to the beads. Each wash was incubated for 30 minutes at room temperature, and the supernatants were collected. Proteins from the flow-through and salt wash fractions were precipitated by acetone precipitation. For final elution, 1× SDS loading buffer was added directly to the beads, and samples were boiled at 95 °C for 5 minutes prior to SDS-PAGE analysis.

### Proteasome purification

To obtain human 26S proteasomes (Ding et al., 2019), human erythrocytes (RBCs purchased from The Israel blood bank) were washed with chilled 1X PBS and lysed in hypotonic buffer (25 mM Tris pH 7.4). The cell debris were separated by centrifugation, and the clear red cell lysate were loaded onto 50 mL DEAE Affigel blue column (Biorad) and washed with Buffer A (25 mM Tris pH 7.4, 10% glycerol, 10 mM MgCl2, 1 mM ATP, 1 mM DTT). Proteasome was eluted with a gradient elution from 0%–50% of Buffer B (25 mM Tris pH 7.4, 10% glycerol, 10 mM MgCl2, 1 mM ATP, 1 mM DTT with 1 M NaCl). The active fractions were then checked by 25 mM Suc-LLVY-AMC activity assay (Glickman & Coux, 2001; Leggett et al., 2005), pooled and loaded onto 8 mL Resource-Q column with Buffer A. Then proteasomes were eluted with a gradient elution from 0%–50% of Buffer B and active fractions were then checked by activity assays conducted either in a 96-well plate or native PAGE. Each active Resource-Q fraction was checked for presence of pDUBs by immunoblotting and process separately for the next step. To obtain the highest purity, gel filtration was performed for each fraction using a S300 (120 mL) column (GE Life Sciences) with Buffer A. The purity and integrity of proteasomes were checked in native-PAGE and SDS-PAGE, then concentrated, aliquoted, flash frozen in liquid nitrogen and stored at −80°C.

### Recombinant Protein purification

Recombinant proteins containing an N-terminal 10×His tag or GST tag were expressed in *E. coli* BL21(DE3) cells. Overnight starter cultures were expanded into 1 L LB medium supplemented with ampicillin (Sigma-Aldrich A9518) and grown at 37 °C to an OD600 of 0.6–0.8. Protein expression was induced with 200 µM IPTG (Inalco 1758-1400), and cultures were incubated overnight at 18 °C. For HIS tag purification: Cells were harvested by centrifugation (5,000 × g, 10 min, 4 °C) and resuspended in HisTrap Buffer A (50 mM Tris pH 7.4, 500 mM NaCl, 20 mM imidazole, 1 mM DTT, 5% glycerol, protease inhibitors). Cells were lysed using a French press, and lysate was clarified by centrifugation (18,000 × g, 30 min, 4 °C) and filtration. The clarified lysate was applied to a HisTrap IMAC column (BioRad 12009300) using NGC system (BioRad), equilibrated with Buffer A. Bound proteins were washed extensively and eluted with a linear imidazole gradient up to 500 mM in Buffer B (composition as Buffer A, with 500 mM imidazole). Peak fractions were analyzed by SDS-PAGE, pooled, and dialyzed overnight at 4 °C in SEC buffer (50 mM Tris pH 7.4, 150 mM NaCl, 1 mM DTT) post dialysis 10% glycerol was added. Dialyzed proteins were concentrated using centrifugal filters (Amicon Ultra Millipore) to ~300 µL and further purified by size-exclusion chromatography (SEC) on a pre-equilibrated column (Cytiva superose 6 10/300 GL 29091596) in SEC buffer. Fractions corresponding to the major peak were pooled, concentrated, flash-frozen in small aliquots, and stored at –80 °C. For GST tag purification: Cells were harvested by centrifugation (5,000 × g, 10 min, 4 °C) and resuspended in lysis/wash Buffer A (50 mM Tris pH 7.4, 150mM NaCl, 1 mM DTT, 5% glycerol, protease inhibitors). Cells were lysed using a French press, and lysate was clarified by centrifugation (18,000 × g, 30 min, 4 °C) and filtration. The clarified lysate was applied to a GSTrap column (Cytiva GE28401748) using NGC system (BioRad), equilibrated with wash buffer. Bound proteins were washed extensively and eluted with a elution buffer (composition as wash buffer, with 20mM GSH). Peak fractions were analyzed by SDS-PAGE, pooled, and dialyzed overnight at 4 °C. Dialyzed proteins were concentrated and further purified by size-exclusion chromatography as described.

### LC-MS/MS and Mass-spectrometry analysis of Proteasome samples

The enriched proteasome samples from different FPLC columns were either processed by in-solution trypsin digestion or native In-gel trypsin digestion, followed by desalting using C18 column. The samples were analyzed using Q-Exactive-Plus or Q-Exactive HF mass spectrometer (*Thermo Fisher*) coupled to nano HPLC. The peptides were resolved by reverse-phase chromatography on 0.075 × 180 mm fused silica capillaries (J&W) packed with Reprosil reversed-phase material (*Dr. Maisch; GmbH, Germany*). The peptides were eluted with a linear 60 min gradient of 5–28%, followed by a 15 min gradient of 28–95%, and a10 min wash at 95% acetonitrile with 0.1% formic acid in water (at flow rates of 0.15 μl/min). Mass spectrometry analysis by Q Exactive HF-X mass spectrometer (*Thermo Fisher Scientific*) was in positive mode using a range of m/z 300–1800, resolution 60,000 for MS1 and 15,000 for MS2, using repetitively full MS scan followed by high energy collisional dissociation (HCD) of the 18 most dominant ions selected from the first MS scan.

MS data analysis was done with either MaxQuant version 1.6.7.0 or the Trans Proteomic Pipeline (TPP) v5.2.0 Flammagenitus using default parameters (Sahu et al., 2021). The raw files were searched against the Homo sapiens UniProt fasta database (November 2017; 20,239 sequences). Candidates were filtered to obtain False discovery rate (FDR) of 1% at the peptide and the protein levels. No filter was applied to the number of peptides per protein. For quantification, the match between runs modules of MaxQuant were used, and the LFQ (label free quantification) normalization method was enabled. For pDUBs abundance calculation in all the proteasome preps the LFQ intensities are normalized with the cumulative/average LFQ values of all 20S proteasome subunits.

### In vitro substrate degradation assay

The substrate degradation assays were performed using 20 nM of human 26S^U15^ or 26S^D-^ proteasomes and ubiquitinated α-globin or Cyclin B1 in a molar ratio of 1:150. The 26S proteasome assay buffer contained 25mM TRIS (pH 7.4), 10mM MgCl2, 10% glycerol, 1 mM ATP, and 1mM DTT. The 20S proteasome assay buffer contained 25mM TRIS (pH 7.4), 150mM NaCl, 10% glycerol without MgCl2 and ATP. All degradation reactions were carried out at 37 °C and were terminated by adding SDS-loading dye.

### AlphaFold3 Models

Models presented in figure 6 were created using the AlphaFold3 package local implementation (https://doi.org/10.1038/s41586-024-07487-w), with 20 different model seeds, such that 100 models were produced in each run.

### Structure analysis and figure preparation

AlphaFold structure predictions and PDB structures were visualized and analyzed using UCSF ChimeraX-1.10 (Pettersen et al., 2021).

### Ranking scores and pyDock energy computations

Binding energies for protein complexes were calculated using the pyDock3 energy scoring function (Jiménez-García et al., 2013), which combines electrostatic, van der Waals, and de-solvation energy terms.

## Author contributions

IS initiated the project. SL and IS performed all biochemical and cell biology assays. ARW and EF aided protein purification. NR designed and supported cloning. SM and SK synthesised synthetic proteins. IB and MJL provided and analysed affinity-tagged proteasomes. FG carried out bioinformatic models and analysis. NC and OK performed MS/MS analysis. MJL, OK, AB, IS, and MHG oversaw the project, supervised, and secured funding. SL, IS, and MHG conceptualized the project and monitored progress. SL, IS, and MHG wrote the first draft of the manuscript. All authors read, commented, and approved the manuscript.

## Acknowledgements and Funding

This work was partially funded by ISF BRG grant 2640/23, and by the European Union ERC grant (UbWan, 101142726) to MHG. Views and opinions expressed are however those of the author(s) only and do not necessarily reflect those of the European Union or the European Research Council Executive Agency. Neither the European Union nor the granting authority can be held responsible for them.

## Data Sharing

All relevant data is included with the submission. The mass spectrometry proteomics data will be deposited to the ProteomeXchange Consortium via the PRIDE partner repository with the Project accession.

