## Supplementary Information for "USP15: the fourth Proteasome-associated DUB"

For manuscript titled:

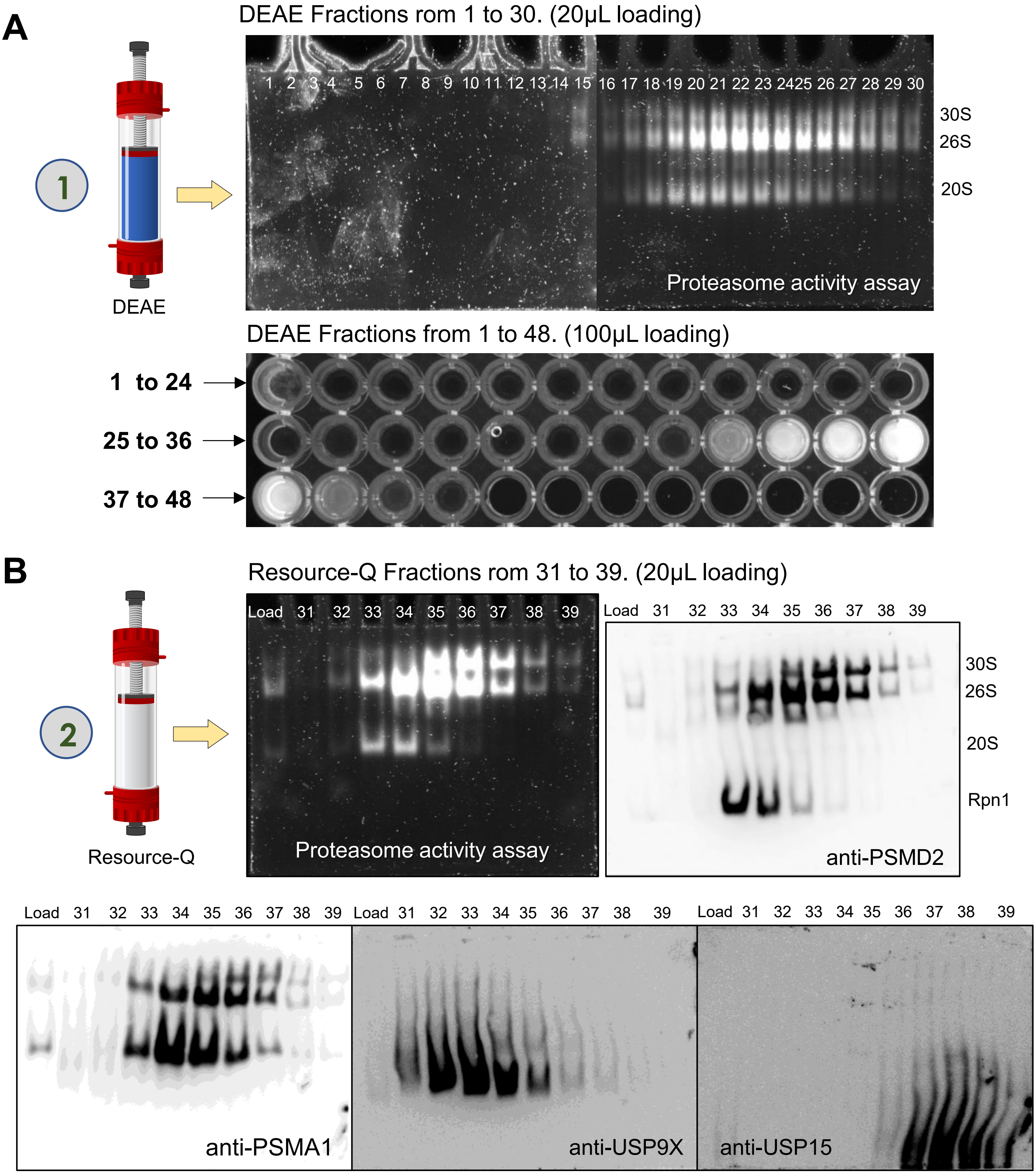


**Sup Figure 1: The 3-step Purification method of 26SU15 proteasome**. (**A**) The RBC lysates were passed through the DEAE affigel blue column and each fractions were run in native gel for in-gel proteasome activity assay (upper) and 96-well plate peptidase assay (lower). The active fractions were considered for the next step of purification. (**B**) The active DEAE fractions are pooled together and passed through the Resource-Q column. The eluted fractions are run in native gels followed by in-gel activity assay and immunoblotting with anti-PSMD2, anti-PSMA1, anti-USP9X, anti-USP15 abs.

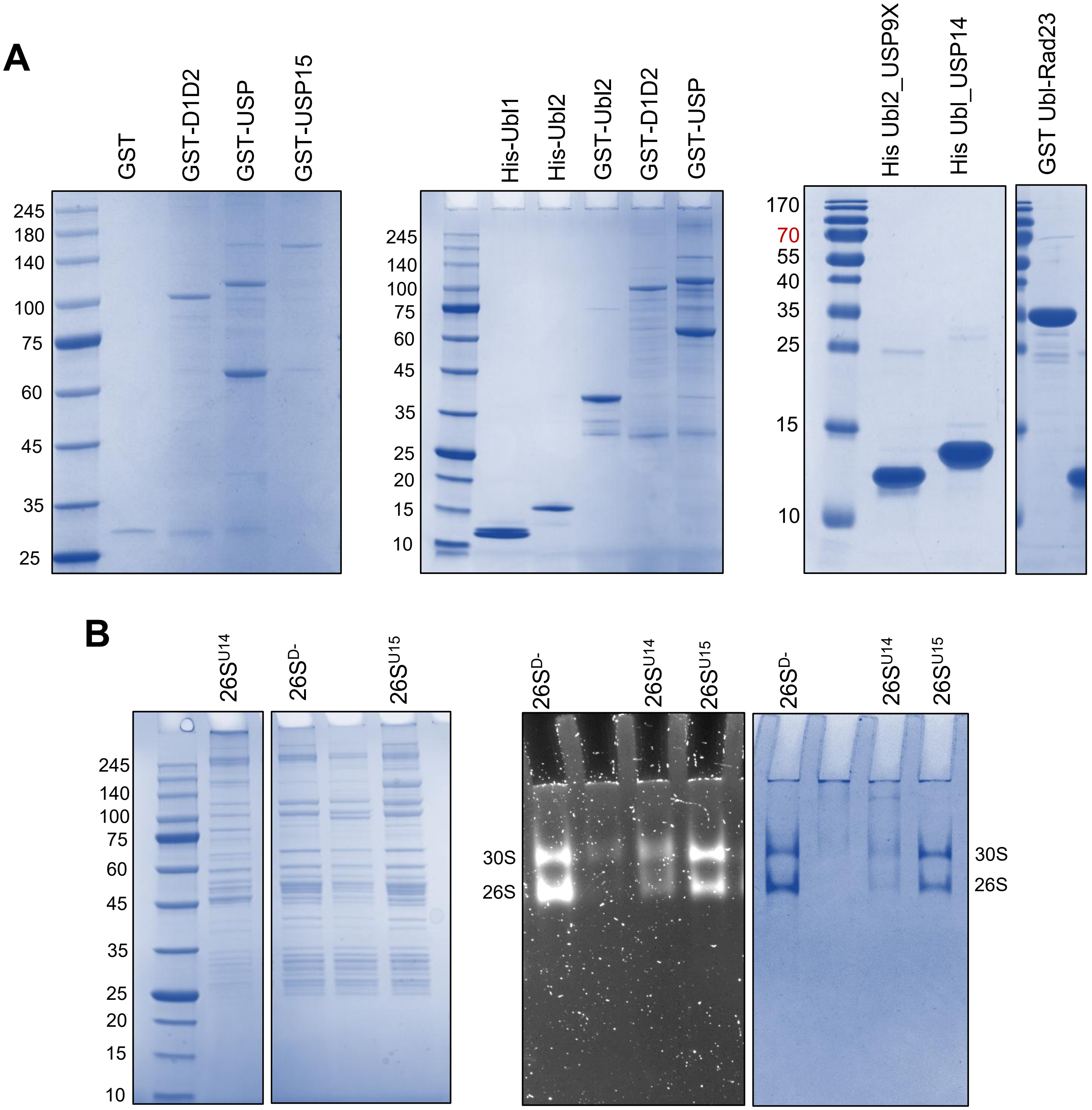


**Sup Figure 2: Purified recombinant proteins and human 26S proteasomes**. (**A**) Recombinant proteins or domains of USP15, USP9X, USP14 and Rad23 were purified either with GST or His tag and run in denaturing-PAGE followed by CBB staining. (**B**) Purified human proteasome ($26S^{U15}$, $26S^{U14}$ and $26S^{D-}$) were run in denaturing gels followed by CBB staining. $26S^{U15}$, $26S^{U14}$ and $26S^{D-}$ proteasomes were run in a 4% native gel followed by In-gel activity assay and CBB staining.

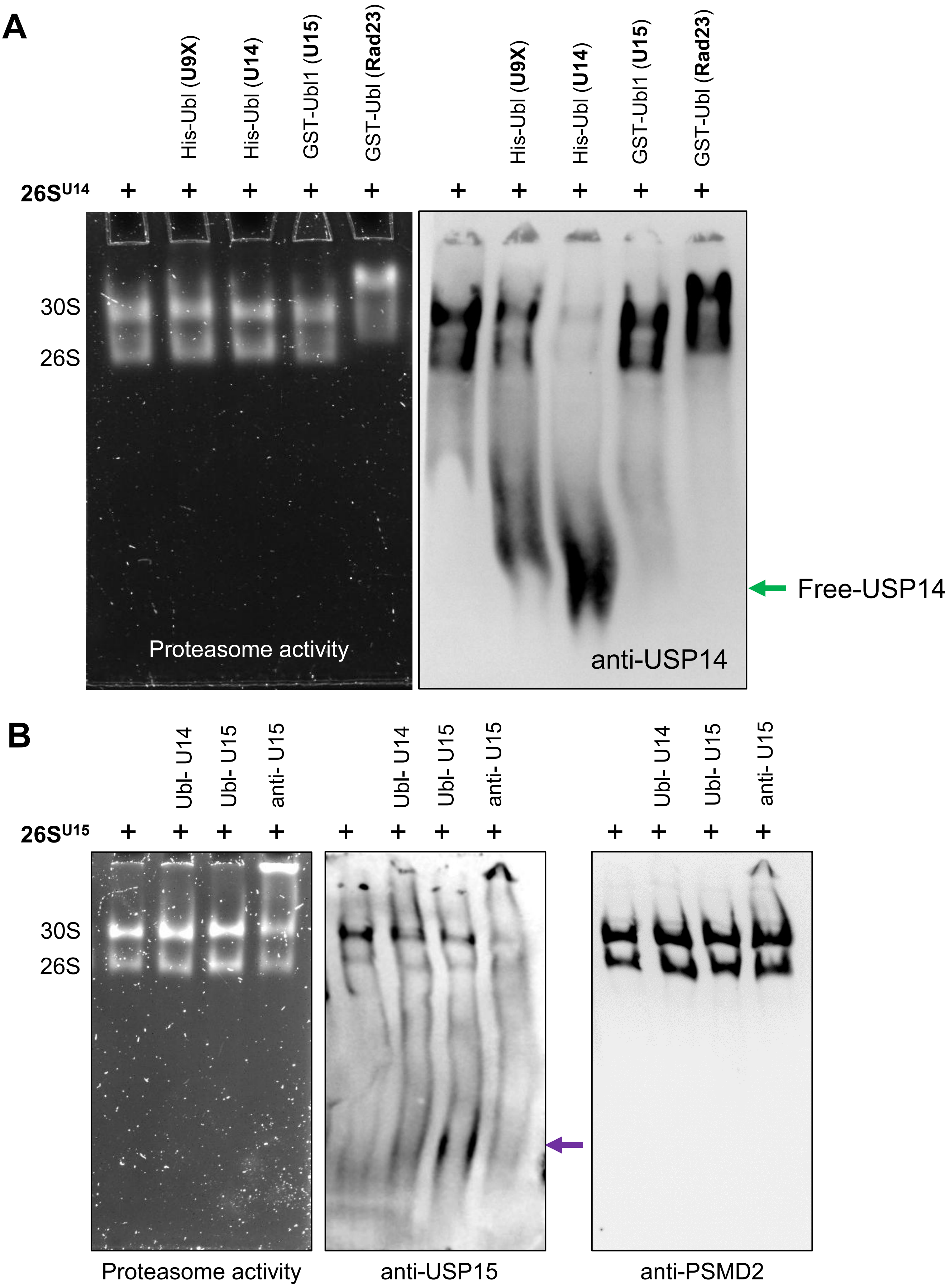
A
His-Ubl (U9X)
His-Ubl (U14)
GST-Ubl1 (U15)
GST-Ubl (Rad23)
26S^U14
30S
26S
Proteasome activity
anti-USP14
Free-USP14
B
Ubl- U14
Ubl- U15
anti- U15
26S^U15
Proteasome activity
anti-USP15
anti-PSMD2

**Sup Figure 3: USP15 does not compete out USP14 on human 26S proteasomes**. (**A**) The $26S^{U14}$ proteasomes were mixed with different Ubl domains as indicated and run in a 4% native gel followed by in-gel activity assays and immunoblotting with anti-USP14 antibody. (**B**) The $26S^{U15}$ proteasomes were mixed with USP15/USP14 Ubl domains or with antiUsp15 antibody as indicated and run in a 4% native gel followed by in-gel activity assays and immunoblotting with anti-USP15 antibody.

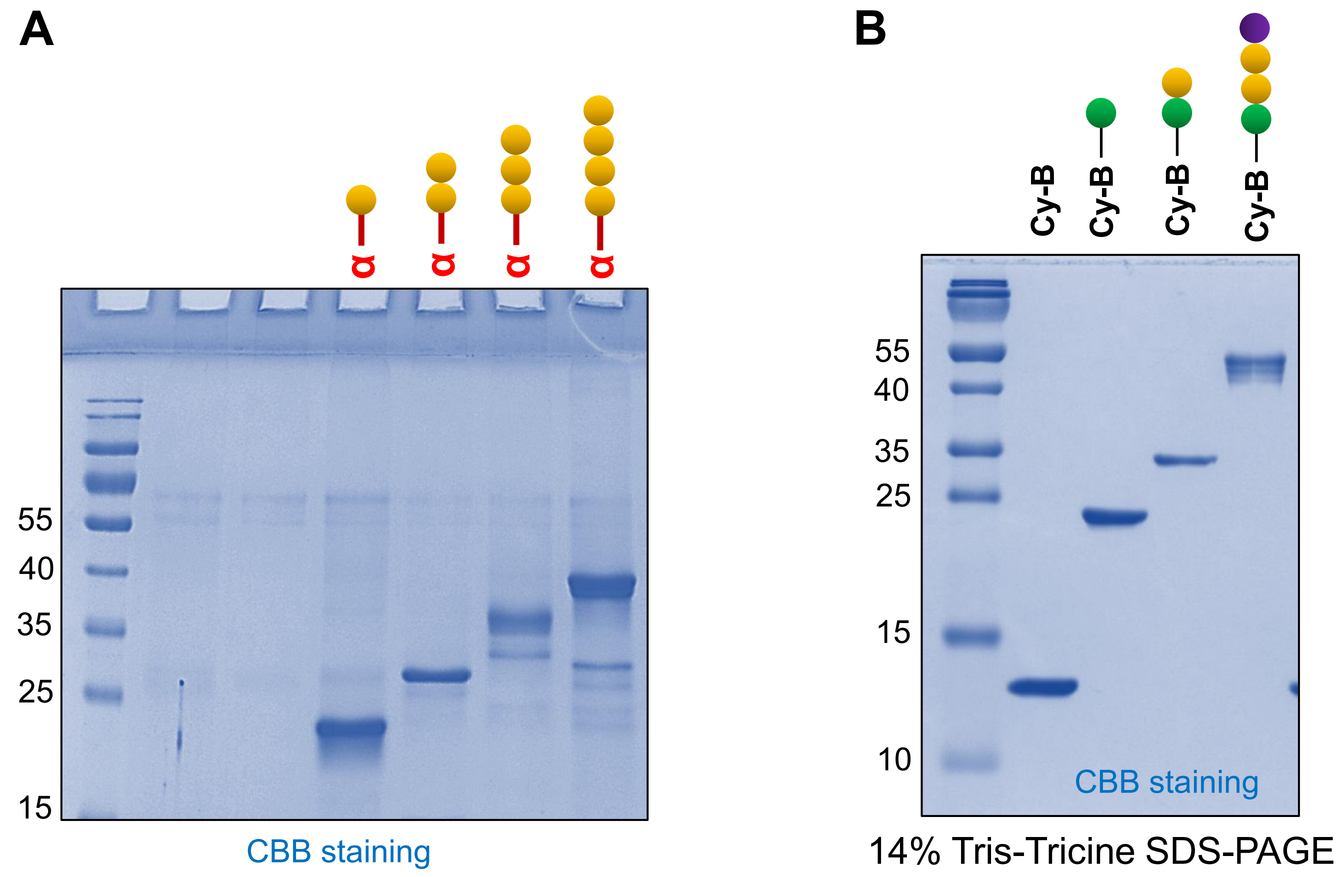


**Sup Figure 4: Purified synthetic and semi-synthetic ubiquitin conjugates.** Chemically synthesized and purified α-globin attached to various units of ubiquitin (**A**) and CyclinB1 attached to various units of tagged ubiquitin (**B**) were run in denaturing PAGE followed by CBB staining.
